# Irgm1-deficiency leads to myeloid dysfunction in colon lamina propria and susceptibility to the intestinal pathogen *Citrobacter rodentium*

**DOI:** 10.1101/575415

**Authors:** Gregory A. Taylor, Hsin-I Huang, Brian E. Fee, Nourhan Yousssef, Viviana Cantillana, Alexi A. Schoenborn, Allison R. Rogala, Anne F. Buckley, Bruce A. Vallance, Ajay S. Gulati, Gianna E. Hammer

## Abstract

*IRGM* and its mouse orthologue *Irgm1* are dynamin-like proteins that regulate vesicular remodeling, intracellular microbial killing, and pathogen immunity. *IRGM* dysfunction is linked to inflammatory bowel disease (IBD), and while it is thought that defective intracellular killing of microbes underscores IBD susceptibility, studies have yet to address how IRGM/Irgm1 regulates immunity to microbes relevant to intestinal inflammation. Here we find that loss of Irgm1 confers marked susceptibility to *Citrobacter rodentium*, a noninvasive intestinal pathogen that models inflammatory responses to intestinal bacteria. Irgm1-deficient mice fail to control *C. rodentium* outgrowth in the intestine, leading to systemic pathogen spread and host mortality. Surprisingly, susceptibility due to loss of Irgm1 function was not linked to defective intracellular killing of *C. rodentium* or exaggerated inflammation, but was instead linked to failure to remodel specific colon lamina propria (C-LP) myeloid cells that expand in response to *C. rodentium* infection and are essential for *C. rodentium* immunity. Defective immune remodeling was most striking in C-LP monocytes, which were successfully recruited to the infected C-LP, but subsequently underwent apoptosis. Apoptotic susceptibility was induced by *C. rodentium* infection and was specific to this setting of pathogen infection, and was not apparent in other settings of intestinal inflammation. These studies reveal a novel role for Irgm1 in host defense and suggest that deficiencies in survival and remodeling of C-LP myeloid cells that control inflammatory intestinal bacteria may underpin IBD pathogenesis linked to IRGM dysfunction.

**Author Summary:** Intestinal macrophages are seeded by peripheral monocytes that enter the intestine and mature into an essential component of immune defense. While this process is shaped by intestinal bacteria, the mechanisms that regulate the process, and their roles in host defense to enteric pathogens are poorly defined. We find that *Irgm1* – the orthologue of the human Crohn’s disease resistance gene, *IRGM* – is required for monocyte/macrophage remodeling in response to the gram-negative enteric pathogen *Citrobacter rodentium*, with absence of Irgm1 leading to systemic spread of the bacterium and host mortality. Failure to remodel colonic monocytes/macrophages was due to a key requirement for Irgm1 in colon-infiltrating monocytes, which underwent dramatically high rates of cell death, specifically during the setting of enteric bacterial infection. While Irgm1/IRGM proteins are well appreciated for their role in regulating intracellular bacterial killing, our results expand the paradigm for Irgm1/IRGM-mediated host defense by demonstrating an essential function for Irgm1/IRGM to control the survival and population remodeling of monocytes and macrophages that expand during enteric infection and defend against pathogenic bacteria. These findings have significant implications for the genesis of Crohn’s disease in individuals that carry *IRGM* variants.

## Introduction

The Immunity Related GTPases (IRG) are a family of large, dynamin-like GTPases that mediate immune and inflammatory responses to pathogenic challenges[1–3]. Their expression is stimulated by interferons and microbial products in both hematopoietic and non-hematopoietic cells, where they bind intracellular membranes and trigger diverse membrane remodeling and vesicle trafficking events. While much about these processes and the mechanisms by which they support immunity remains unclear, the importance of IRG functions are underscored by the existence of variants in the human *IRGM* gene that are associated with increased susceptibility to *Mycobacterium tuberculosis* [4, 5], sepsis[6], non-alcoholic fatty liver disease[7], ankylosing spondylitis[8], and Crohn’s Disease (CD). CD is an inflammatory bowel disease thought to result from aberrant immune responses to intestinal microbes and CD-linked *IRGM* variants are linked with the overall incidence of CD[9, 10] but also with disease severity including ileal involvement[11], fistulating disease [12], and an increased need for surgery[13]. *IRGM* is the only IRG that is widely expressed in human cells and despite strong genetic evidence linking *IRGM* variants with CD, the means through which *IRGM* dysfunction confers CD susceptibility remains unknown.

Functional studies have demarcated a role for IRGM and its mouse orthologue, *Irgm1*, in regulating autophagy [14–18] and bacterial outgrowth during pathogenic infection. Irgm1-deficient mice are highly susceptible to intracellular bacteria including *Listeria monocytogenes[19]*, *Salmonella typhimurium[20]*, and *Mycobacterium tuberculosis*[21]. Susceptibility to these pathogens is linked to impaired abilities of Irgm1-deficient macrophages to kill intracellular bacteria [20, 21], a functional deficiency that is also observed in IRGM-deficient human macrophages [15, 22]. Based on these studies, the prevailing model by which IRGM/Irgm1 controls immunity is by supporting mechanisms of intracellular microbial killing, including xenophagy. It is speculated that impaired xenophagy and microbial killing impacts immune responses to intestinal microbes relevant to CD. However, this model rests largely on *in vitro* studies employing cultured macrophages and little information exists concerning how IRGM/Irgm1 regulates macrophages and other immune cells *in vivo*.

Given the evidence linking IRGM/Irgm1 to both bacterial immunity and aberrant immunity in CD[17, 23], it is surprising that no study has addressed how IRGM/Irgm1 regulates the functions of intestine-resident immune cells, or host immunity to intestinal pathogens. Moreover, with the focus of previous work on invasive, intracellular bacteria, little is known about IRGM/Irgm1’s potential roles in promoting resistance to extracellular bacteria, by far the most abundant type of microbe in the intestine and key players in intestinal inflammation. A model organism in this class is *Citrobacter rodentium*, a Gram-negative, enteric pathogen of mice that induces intestinal inflammation sharing several histologic features with IBD [24, 25]. *C. rodentium* is noninvasive, and its infection resembles human infection with attaching and effacing *E. coli*[26], clinically important microbes that cause diarrhea and gastrointestinal inflammation. Numerous studies have shown that *C. rodentium* infection is typically self-resolving, due to robust responses of both innate and adaptive immune cells[24, 25]. Here we find that Irgm1 plays an important role in protective immunity to *C. rodentium*, but one that is distinct from its role in xenophagy and intracellular killing.

## Results

### Irgm1 function is required for *C. rodentium* immunity

We inoculated mice that have a global knockout of Irgm1 (*Irgm1*^−/−^ mice)[19] with *C. rodentium* intragastrically, and followed the course of infection over three weeks. While the vast majority of wild-type (WT) mice survived *C. rodentium* infection, ~40% of *Irgm1*^−/−^ mice died between 12 and 20 days post-infection (Fig 1a). During that period, *Irgm1*^−/−^ mice also manifested marked weight loss that was not apparent in the control mice (Fig 1b). To determine if Irgm1 function controlled *C. rodentium* outgrowth in the intestine, we quantified the abundance of *C. rodentium* in stool at day 7, 14 and 21 post-infection. Surprisingly, at day 7 the bacterial load was not significantly different between WT and *Irgm1*^−/−^ mice (Fig 1c). However, while WT mice restricted pathogen outgrowth and sharply reduced bacterial load within 14 days post infection, *C. rodentium* continued to expand in the intestines of *Irgm1*^−/−^ mice, resulting in a bacterial burden 20-30-fold higher than that of WT mice. By day 21 many *Irgm1*^−/−^ mice had succumbed to the infection, and those that survived still had detectable levels of bacteria in stool. Long-term persistence of *C. rodentium* in *Irgm1*^−/−^ mice was in marked contrast to WT, in which no bacteria were detected at late timepoints. Taken together, these results suggest that Irgm1 function was required for immunity to *C. rodentium* and to restrict pathogen outgrowth in the intestine.

**Figure 1.**
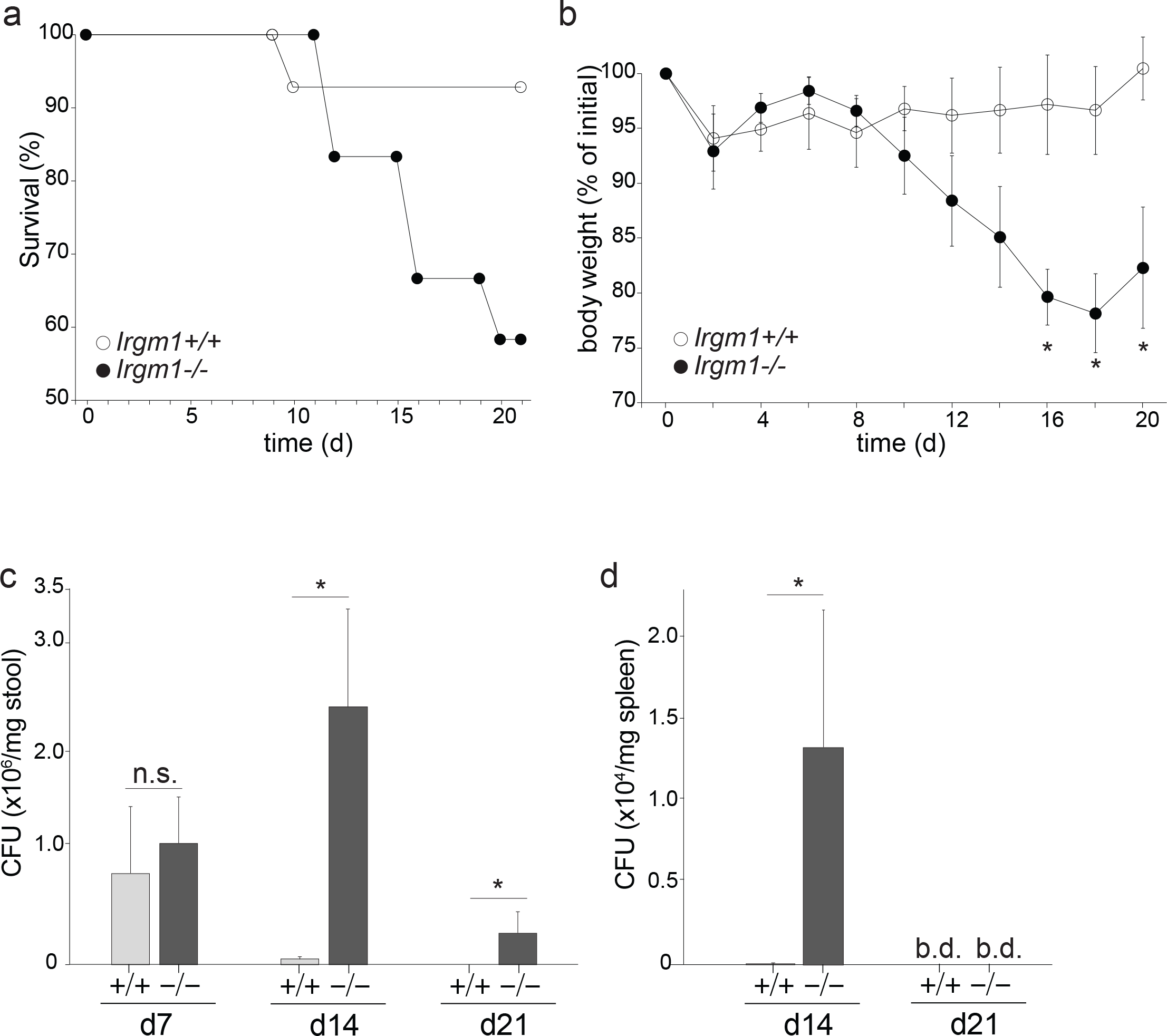
Irgm1 is required for resistance to the enteric pathogen *C. rodentium*. *Irgm1*^+/+^ and *Irgm1*^−/−^ mice were infected with *C. rodentium* and evaluated for survival (a) and weight loss (b) over the period of 21 days. The abundance of *C. rodentium* in stool (c) or spleen (d) of infected mice was evaluated in tandem. (a) n=12 *Irgm1*^+/+^ mice and n=11 *Irgm1*^−/−^ mice; *p* =0.05, Log Rank (Mantel-Cox) test. (b) n=6. (c) n=6. (d) n=5. (a) - (c) \**p* < 0.05, unpaired Student’s t test; n.s., not significant; b.d., below detection; error bars indicate SEM. The data are representative of 3 experiments.

Given these results, we assessed whether failure to control *C. rodentium* in the intestine of *Irgm1*^−/−^ mice allowed bacteria to spread systemically. *C. rodentium* is non-invasive and its systemic spread is both rare and limited to hosts with significant defects in intestinal barrier function (e.g. [27, 28]). Consistent with this model, we detected no *C. rodentium* in the spleens of infected WT mice (Fig 1d). By contrast, several thousand live colony forming units of *C. rodentium* were detected in the spleens of *Irgm1*^−/−^ mice, and this abundance peaked at day 14 post-infection, a timepoint where exaggerated weight loss and mortality of *Irgm1*^−/−^ mice became evident. These data suggest that Irgm1 function is required to restrict pathogen expansion in the intestine, as well as its systemic spread.

### Dysfunctional *C. Rodentium* immunity in *Irgm1*-deficient mice is not linked to exaggerated intestinal inflammation

*C. rodentium* reproducibly causes intestinal inflammation and is a standard model to study host dynamics in infectious colitis. *Irgm1*^−/−^ mice develop robust colitis upon treatment with dextran sodium sulfate (DSS colitis)[17] and given that *C. rodentium* infected *Irgm1*^−/−^ mice poorly controlled pathogen outgrowth, displayed heightened mortality and exaggerated weight loss, we hypothesized that infected *Irgm1*^−/−^ mice would also display exaggerated intestinal inflammation and colitis. We focused on day 14 post infection, as it was the point at which *Irgm1*^−/−^ mice exhibited clear signs of mortality and weight loss (Fig 1). In contrast to our prediction, inflammatory scores were not significantly different between *Irgm1*^−/−^ and WT mice (Fig 2a-d). We performed extensive histologic analysis for edema, epithelial hyperplasia, epithelial integrity, and cellular infiltrate in both cecum and distal colon, the primary and secondary sites for *C. rodentium* replication. In the cecum, none of the 4 parameters analyzed nor the aggregate inflammation score were statistically different between *Irgm1*^−/−^ and WT mice (Fig 2a,b). Only edema in distal colon showed a statistically significant increase in *Irgm1*^−/−^ mice, although this difference was modest and did not significantly impact the aggregate inflammation score (Fig 2c,d). Pathology was scored in a blinded fashion by three independent individuals, with similar results (data not shown). As an additional measure for inflammation we quantified gene expression for TNFα in colon tissues. TNFα mRNA was upregulated in colon following infection, but levels in WT and *Irgm1*^−/−^ mice were very similar, even at time points when mortality and weight loss of *Irgm1*^−/−^ mice were readily apparent (Fig 2e). Taken together, these data suggest that mechanisms underpinning defective *C. rodentium* immunity in *Irgm1*^−/−^ mice and pathogen outgrowth were not linked to the severity of intestinal inflammation.

**Figure 2.**
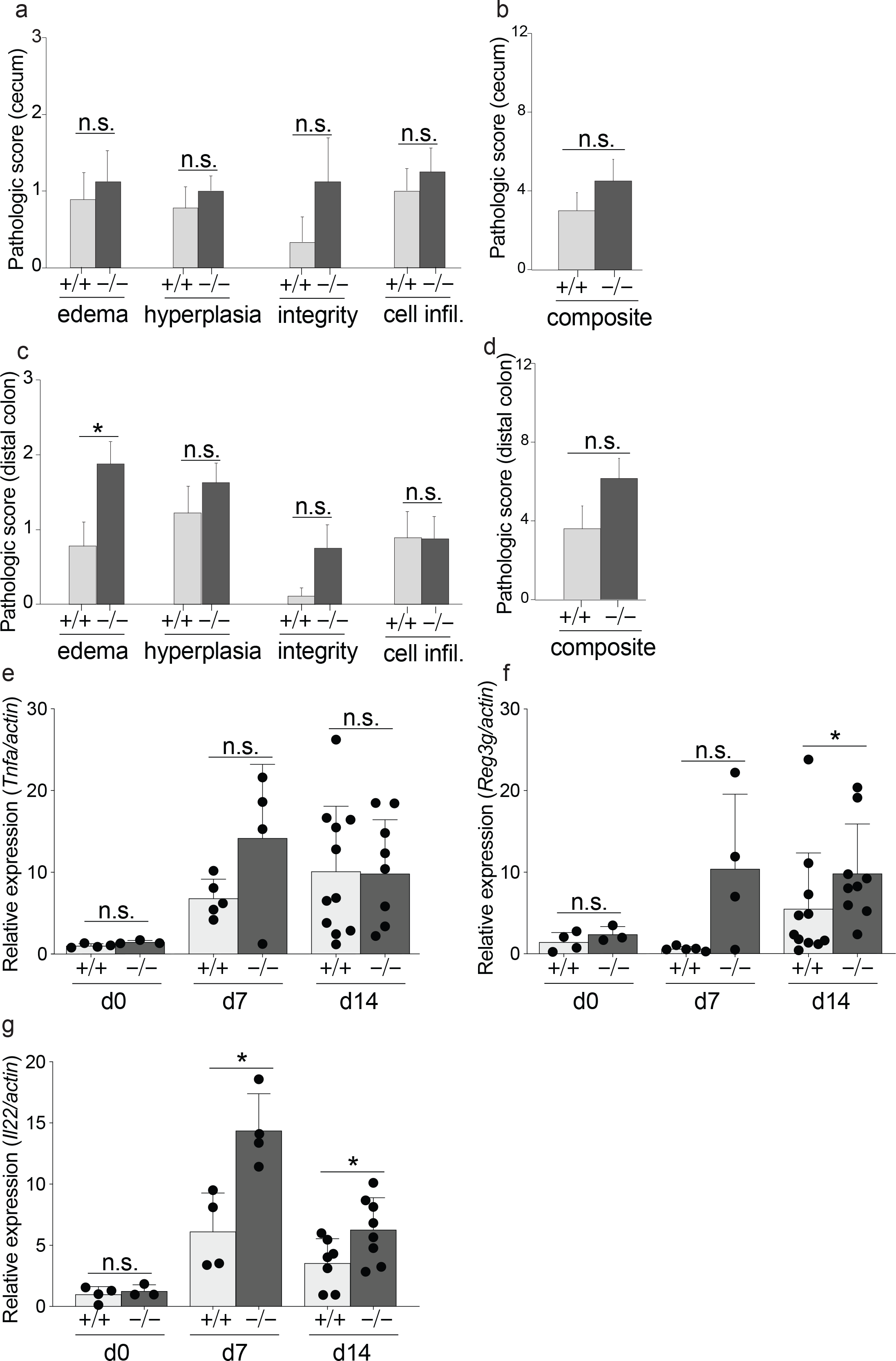
Susceptibility of *Irgm1*^−/−^ mice to *C. rodentium* is not linked to exaggerated intestinal inflammation. Cecum from *Irgm1*^+/+^ and *Irgm1*^−/−^ mice at 10 days post-infection with *C. rodentium* were analyzed for the indicated inflammatory pathology (a). The composite of all inflammatory pathologies in cecum is shown in (b). (c,d) Distal colon was analyzed as above. Separate cohorts of *C. rodentium* infected mice were analyzed for the abundance of mRNA expression of TNFα (e), Reg3Ɣ (f), and IL-22 (g) in distal colon. Data were combined from two independent experiments. (a-d) n=8-9, error bars indicate SEM. (e-f) Each dot represents one mouse, error bars indicate SD. \**p* < 0.05, n.s., not significant (unpaired Student’s t test).

To further investigate the inflammatory response against *C. rodentium*, we quantified expression of *Reg3γ*, an antimicrobial peptide produced by intestinal epithelial cells that suppresses *C. rodentium* outgrowth and is known to be expressed in an Irgm1-dependent fashion during DSS colitis[17, 29]. In response to *C. rodentium* infection *Reg3γ* was robustly expressed in colon tissue, and expression was actually elevated in *Irgm1*^−/−^ mice, perhaps due to exaggerated pathogen expansion in these mice (Fig 2f). *C. rodentium* infected *Irgm1*^−/−^ mice also displayed robust and elevated gene expression of IL-22, an essential cytokine for host defense against *C. rodentium* and tissue repair in the intestine[30] (Fig 2g). Taken together, these histologic and gene expression data suggest that *Irgm1*^−/−^ mice elicit an apparently normal tissue inflammatory and repair response to *C. rodentium* and that pathogen outgrowth, systemic spread, and mortality cannot be explained by “general susceptibility” of *Irgm1*^−/−^ mice to intestinal inflammation, or by deficiency in key host defense cytokines.

### Loss of Irgm1 function compromises immune remodeling of monocyte/macrophage populations in response to *C. rodentium*

Pathogens and other inflammatory stimuli induce *Irgm1* expression in both immune and non-immune populations and there are thus multiple potential sources in which Irgm1 could be critical for host defense to *C. rodentium*. Since the cellular responses of intestinal epithelial cells to *C. rodentium* appeared to be independent of Irgm1, we hypothesized that mechanisms underpinning dysfunctional *C. rodentium* immunity in *Irgm1*^−/−^ mice were linked to hematopoietic cells. To test this we generated *Irgm1*^−/−^ hematopoietic chimeric mice by adoptively transferring *Irgm1*^−/−^ bone marrow into irradiated WT recipients. *Irgm1*^−/−^ → WT hematopoietic chimera were highly susceptible to *C. rodentium* infection and manifested severe weight loss and early mortality, paralleling outcomes with *Irgm1*^−/−^ mice (Fig S1). Thus, *Irgm1*-deficiency in hematopoietic cells is sufficient for dysfunctional *C. rodentium* immunity.

Having identified hematopoietic cells as the source of *C. rodentium* susceptibility in *Irgm1*^−/−^ mice we investigated macrophages, since these are the most abundant phagocytes in the intestine, and multiple lines of evidence support Irgm1 as a critical regulator of macrophage-intrinsic mechanisms of pathogen defense[20, 21]. Using *Irgm1*^−/−^ bone marrow derived macrophages we tested whether Irgm1 was required for macrophages to uptake and kill intracellular *C. rodentium*, and surprisingly found that these defense mechanisms were both robust and undiminished as compared to WT (Fig 3). This outcome is in contrast to prior reports showing a requirement for Irgm1 for macrophage killing of *Salmonella typhimurium*[20] and *Mycobacterium tuberculosis*[21]. Thus, deficiency in *C. rodentium* killing by macrophages does not explain why *Irgm1*^−/−^ mice fail to control *C. rodentium* infection.

**Figure 3.**
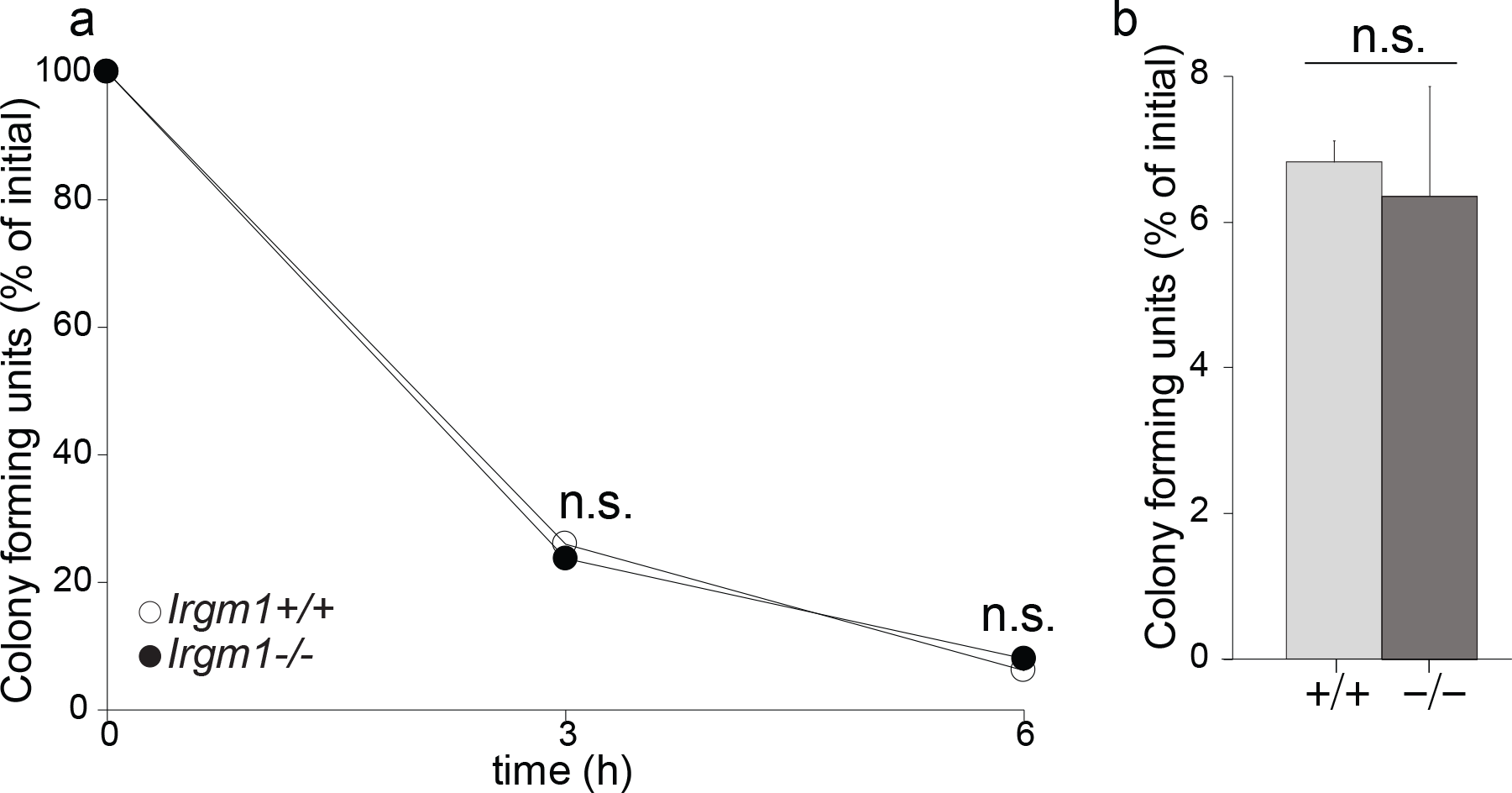
Macrophage killing of *C. rodentiu*m is independent of Irgm1 function. Bone marrow macrophages of *Irgm1*^+/+^ and *Irgm1*^−/−^ mice were incubated with *C. rodentium* and the killing of internalized bacteria was quantified over time. Data shown is the outcome of a representative experiment (a) and the average of data combined from four independent experiments at the 6h time point (b). Each experiment included bone marrow macrophages from 3 mice per genotype, which were each analyzed independently. Error bars indicate SEM. n.s., not significant (unpaired Student’s t test). Abbreviations: cell infil., neutrophil and mononuclear cellular infiltration.

To specifically investigate Irgm1’s function in colon macrophages and other immune populations in the intestinal mucosa, we extracted cells from the colonic lamina propria (C-LP) and analyzed them by flow cytometry. Unlike most tissues, the majority of C-LP macrophages express high levels of MHC-II (population III, Fig 4a and FigS2) and are steadily replenished by blood-derived monocytes (Ly6C^+^MHC-II^−^; population I), which upregulate MHC II in the intestine (transitioning monocytes, Ly6C^+^MHC-II^+^; population II)[31–33]. Consistent with previous reports, we also found a population of macrophages that expressed low levels of MHC II (population IV)[31]. Whether MHC-II^lo^ macrophages derive from MHC-II^hi^ macrophages or have distinct origins is currently unclear, and since MHC-II expression clearly distinguished two macrophage populations, we analyzed each independently. At steady-state the numbers of C-LP MHC-II^lo^ macrophages, MHC-II^hi^ macrophages, and their monocyte precursors were very similar between *Irgm1*^−/−^ and WT mice (Fig 4c-e and Fig S3). *C. rodentium* infection is known to induce significant remodeling of monocytes/macrophages, and while these populations were indeed profoundly altered after infection, infection-induced remodeling was markedly distinct between *Irgm1*^−/−^ and WT mice (Fig 4a,b). First, WT mice numerically expanded mature MHC-II^hi^ macrophages 2-fold upon *C. rodentium* infection (Fig 4c). By contrast, macrophage expansion did not occur in *C. rodentium* infected *Irgm1*^−/−^ mice, suggesting that these mice were deficient in mechanisms that support remodeling and expansion of C-LP macrophages in response to *C. rodentium*. Because mature macrophages are not proliferative, failure to expand macrophages in infected *Irgm1*^−/−^ mice suggests underlying defects in C-LP macrophage precursors. Consistent with this hypothesis, infected *Irgm1*^−/−^ mice were markedly deficient in Ly6C^+^MHC-II^+^ transitioning monocytes, a direct precursor of C-LP macrophages and a population known to be key for *C. rodentium* immunity (Fig 4d)[31, 34]. Thus, while the WT response to *C. rodentium* elicited >15-fold expansion of Ly6C^+^MHC-II^+^ transitioning monocytes, this population underwent little to no expansion in *Irgm1*^−/−^ mice, and total numbers were numerically similar to that of uninfected controls (Fig 4d). Failure to expand transitioning monocytes in *Irgm1*^−/−^ mice was not due to a deficiency in monocyte precursors, since these expanded 4-fold after infection and were actually increased in abundance as compared to WT (Fig 4e). Exaggerated numbers of monocytes, yet deficiency in their progeny, suggest that in response to *C. rodentium* infection the developmental transition of monocytes into C-LP macrophages was an Irgm1-dependent process.

**Figure 4.**
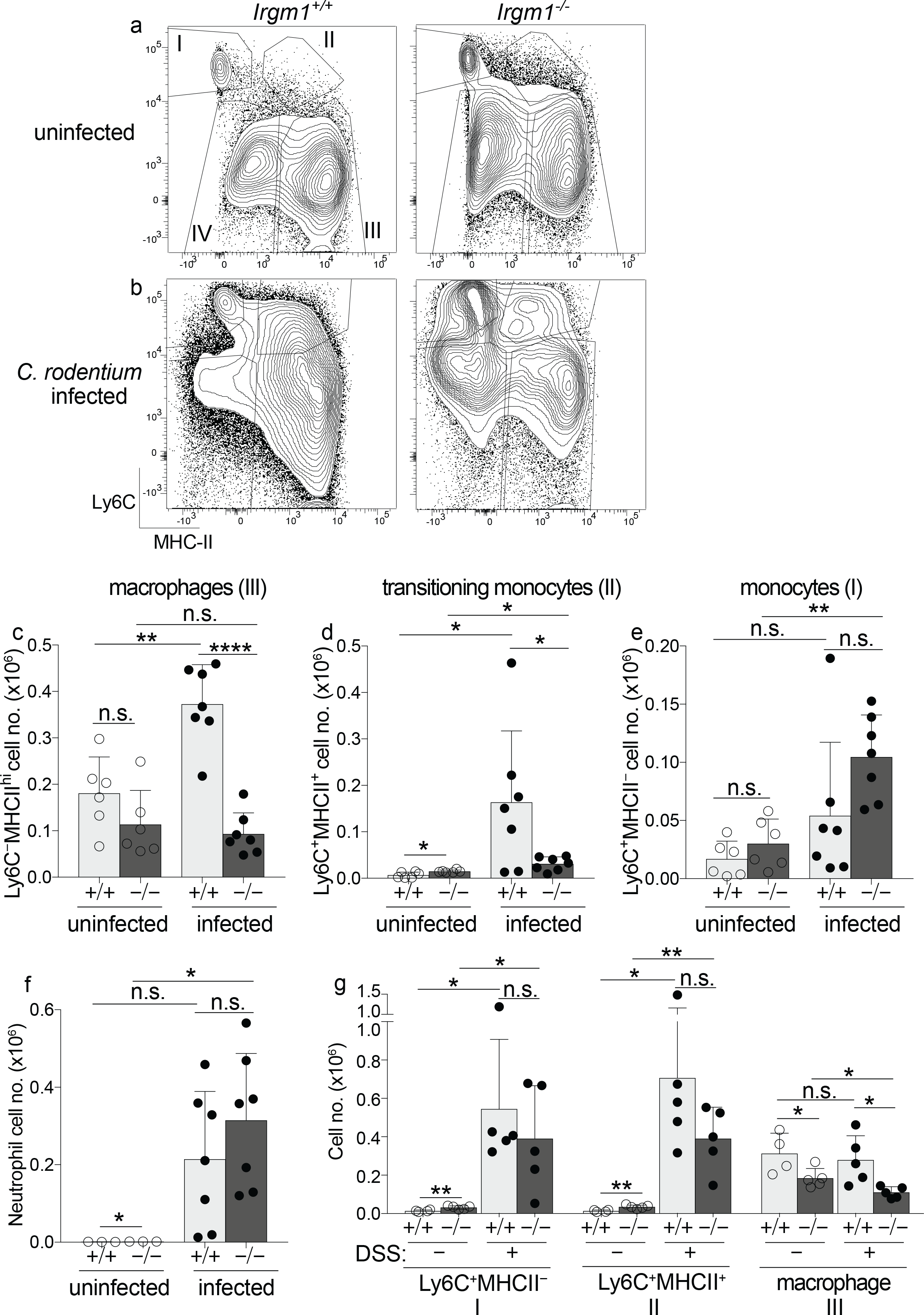
Irgm1 is required for immune remodeling of C-LP monocytes and macrophages in response to *C. rodentium.* Representative flow plots gated on C-LP cells of the monocyte/macrophage lineage from uninfected (a) and day 10 of *C. rodentium* infected (b) *Irgm1*^+/+^ and *Irgm1*^−/−^ mice. C-LP monocytes (I. Ly6C^+^MHCII^−^), transitioning monocytes (II. Ly6C^+^MHCII^+^), and macrophages (III. Ly6C^−^MHCII^hi^, IV. Ly6C^−^MHC-II^lo^) were gated as described in Fig. S2. The total cell number of C-LP monocytes (c), transitioning monocytes (d), Ly6C^−^MHCII^hi^ macrophages (e), and neutrophils (f) in uninfected and *C. rodentium* infected mice of the indicated genotype. (g) C-LP cells of the monocyte/macrophage lineage were quantified from co-housed *Irgm1*^+/+^ and *Irgm1*^−/−^ littermates treated with DSS for six days. Mice given no DSS were Analyzed in tandem. Each dot represents one mouse. Data in (c-f) were combined from two independent experiments using co-housed littermates for each experimental condition. Error bars represent mean ± SD. \**p* < 0.05, \*\**p* < 0.01, \*\*\*\**p* < 0.0001, n.s., not significant (unpaired Student’s t test).

Failure to remodel monocyte/macrophage populations was not due to immunological “ignorance” to *C. rodentium* or general myeloid failure since neutrophil expansion, a hallmark of *C. rodentium* immunity, was robust and increased >150-fold over baseline in infected *Irgm1*^−/−^ mice (Fig 4f). That neutrophil expansion was intact in *Irgm1*^−/−^ mice further suggested that only select myeloid cell subsets had a critical requirement for Irgm1 to undergo remodeling for *C. rodentium* immunity. In further support of this we found no requirement for Irgm1 in MHC-II^lo^ macrophages (Fig S3), suggesting that a critical role for Irgm1 function for *C. rodentium* immunity was cell type specific, even among cells of the monocyte/macrophage lineage. Of note however, MHC-II^lo^ macrophages did not expand or undergo significant remodeling in response to *C. rodentium* infection, which is in marked contrast to the changes that occurred in monocytes/transitioning monocytes and MHC-II^hi^ macrophages. Since remodeling of the latter cell types was Irgm1-dependent, these results may suggest that cellular requirements for Irgm1 function were most critical in those monocyte/macrophage populations which expand and/or differentiate in response to *C. rodentium* infection.

Given these results we tested whether Irgm1 also regulated monocyte/macrophage remodeling during DSS colitis, an inflammatory condition driven by gut microbiota and intestinal barrier defects that induce monocyte influx and monocyte/macrophage remodeling, similar to what occurs during *C. rodentium* infection. In contrast to the monocyte/macrophage remodeling defects of *Irgm1*^−/−^ mice in response to *C. rodentium*, monocytes and transitioning monocytes were robustly expanded during DSS colitis and their numbers were not significantly different between mice of either genotype (Fig 4g). DSS treated *Irgm1*^−/−^ mice did have a reduced number of macrophages, but unlike the WT response to *C. rodentium*, the number of macrophages in WT mice did not increase during DSS colitis, suggesting that the cause for macrophage decline in *Irgm1*^−/−^ mice in DSS colitis was distinct from mechanisms underpinning failure to expand macrophages for *C. rodentium* immunity. Taken together, these data highlight stimulus-specific immune remodeling in C-LP and suggest that Irgm1’s function in monocyte/macrophage remodeling is context dependent, and especially important upon intestinal infection with *C. rodentium*.

### Irgm1 function is required to support CD4 T cell IFNγ responses to *C. rodentium* infection

Despite marked deficiencies in monocyte/macrophage remodeling, all *Irgm1*^−/−^ mice survived to at least 10 days post infection, a time point at which adaptive immunity would be expected to play a role in host defense. Since monocyte/macrophage deficiencies are linked to deficiencies in *C. rodentium*-induced IFNγ-producing CD4 T cells (Th1) [34], we hypothesized that these would be deficient in *Irgm1*^−/−^ mice. Indeed, expansion of Th1 cells following *C. rodentium* infection was significantly impaired, reduced to just 30% that of WT (Fig 5a-c). By contrast, numbers of IL-17-producing CD4 T cells (Th17) and Foxp3^+^ T regulatory cells were intact (Fig 5d and S4), suggesting that among CD4 T cells Irgm1 was specifically required for expansion of IFNγ-mediated host defense against *C. rodentium*. Consistent with this requirement, IFNγ^+^IL-17^+^ CD4 T cells were also modestly reduced (Fig 5e). In further agreement, gene expression for IL-12b (p40), which in the form of IL-12 supports Th1 responses, was markedly deficient in colon tissue of *C. rodentium* infected *Irgm1*^−/−^ mice, and there was also a trend towards decreased gene expression of IL-12a (p35), although the latter did not reach significance (Fig. S5). In contrast to IL-12, *Irgm1*^−/−^ mice had robust gene expression for IL-23 (p19) and IL-1β, cytokines traditionally linked to Th17 responses, further supporting the conclusion that Irgm1’s role in adaptive immunity to *C. rodentium* was most important for IFNγ-mediated CD4 T cell responses.

**Figure 5.**
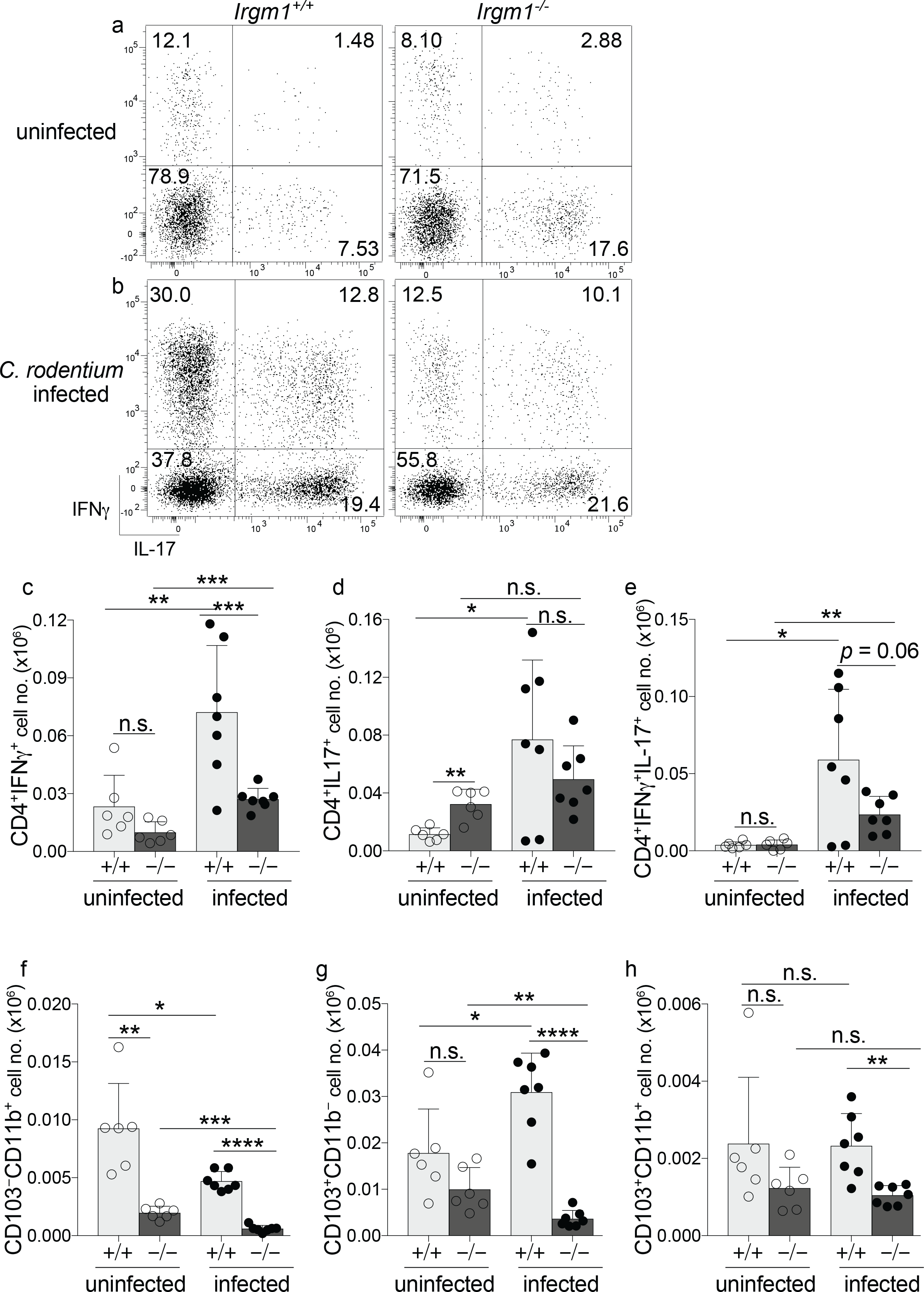
Irgm1 is required for IFNγ-producing CD4 T cells and resilience of dendritic cells in host defense against *C. rodentium*. Representative flow plots of PMA/ionomycin stimulated C-LP CD4 T cells from uninfected (a) and day 10 of *C. rodentium* infected (b) *Irgm1*^+/+^ and *Irgm1*^−/−^ mice. Total cell number of C-LP CD4 T cells producing IFNγ (c), IL-17 (d), or IFNγ^+^IL-17^+^(d) in uninfected or day 10 of *C. rodentium* infected mice of the indicated genotype. These same mice were also analyzed for the total cell number of C-LP CD103^−^CD11b^+^ DCs (f), CD103^+^CD11b^−^ DCs (g), and CD103^+^CD11b^+^ DCs (h). Data were combined from three independent experiments using co-housed littermates for each experimental condition. Each dot represents one mouse. Error bars represent mean ± SD. \**p* < 0.05, \*\**p* < 0.01, \*\*\**p* < 0.001, \*\*\*\**p* < 0.0001, n.s., not significant (unpaired Student’s t test).

Given the defects in IFNγ-producing CD4 T cells, we also analyzed dendritic cells (DCs), since these are required to activate naïve T cells and induce protective *C. rodentium* immunity[35–37]. There are three DC subsets in colon and all three were significantly reduced in *C. rodentium* infected *Irgm1*^−/−^ mice (Fig 5f-h). The subsets most significantly affected were CD103^+^CD11b^−^ and CD103^−^CD11b^+^ DCs, both of which instruct intestinal Th1 responses[38–41]. CD103^−^CD11b^+^ DCs were unique in that their abundance was profoundly reduced in both infected and uninfected *Irgm1*^−/−^ mice (Fig, 6f). CD103^+^CD11b^−^ DCs, the dominant APC subset for intestinal Th1 cells at steady-state[40, 41], were of normal abundance in uninfected *Irgm1*^−/−^ mice, but were markedly reduced 8-fold upon *C. rodentium* infection (Fig 5g). Since *Irgm1*^−/−^ mice had normal numbers of Th1 cells during steady-state (Fig 5c), these data suggest Irgm1’s function to support DCs and/or other antigen presenting cells (APCs) for Th1 responses was most critical for Th1 cells expanded for *C. rodentium* immunity. Taken together our data suggest that marked deficiencies in C-LP DCs and APCs derived from monocyte/macrophage remodeling in *Irgm1*^−/−^ mice impair Th1 responses for *C. rodentium* immunity. The combined defects in innate and adaptive immune responses in infected *Irgm1*^−/−^ mice can be reasonably proposed as the cause for *C. rodentium* outgrowth and systemic spread.

### Myeloid cell deficits in C-LP of Irgm1-deficient mice is underscored by increased apoptotic cell death following *C. rodentium* infection

Given that *Irgm1*^−/−^ mice had defects in innate and adaptive immune responses to *C. rodentium* infection, we sought to identify the cell-intrinsic requirements for Irgm1 in myeloid or lymphoid cells. To do so, we generated mixed hematopoietic chimeric mice by reconstituting WT mice that expressed both CD45.1 and CD45.2 (CD45.1^+^CD45.2^+^) with a mixture of cells from both WT (CD45.1) and *Irgm1*^−/−^ (CD45.2) bone marrow. In a mixed setting *Irgm1*^−/−^ bone marrow poorly competes for immune niche reconstitution[42] and although we injected WT and *Irgm1*^−/−^ bone marrow at a ratio of 1:5, eight weeks post reconstitution C-LP myeloid cells originating from WT bone marrow outnumbered *Irgm1*^−/−^ cells 8:1, with many *Irgm1*^−/−^ myeloid cell types present at even lower frequencies (Fig 6a,b, and Fig. S6). Because the majority of cells in mixed hematopoietic chimera were of wild-type origin we expected these mice to have effective immunity against *C. rodentium*, and we leveraged this phenotype to determine whether the defective cellular responses of Irgm1-deficient cells were independent from the overall failure to control *C. rodentium* outgrowth in *Irgm1*^−/−^ mice. We infected mixed hematopoietic chimera with *C. rodentium* and used CD45.1 and CD45.2 antibodies to distinguish WT and *Irgm1*^−/−^ cells present in the same host. Unlike myeloid cells, CD4 T cells originating from WT or *Irgm1*^−/−^ bone marrow were of equal abundance after *C. rodentium* infection and strikingly, the IFNγ response of *Irgm1*^−/−^ CD4 T cells was identical to that of WT cells in the same host (Fig 6c and Fig. S6). This result contrasts with that of *Irgm1*^−/−^ mice, which were markedly deficient in *C. rodentium*-induced Th1 responses (Fig 5c). Taken together, these data suggest that IFNγ responses to *C. rodentium* did not require Irgm1 function in CD4 T cells themselves, but instead required Irgm1 function in other hematopoietic cells. We thus analyzed cell-intrinsic requirements for Irgm1 in myeloid cells, but found this analysis challenging since unlike CD4 T cells, the ratio of WT: *Irgm1*^−/−^ cells among monocytes, macrophages and DCs dropped sharply after *C. rodentium* infection, and also became highly variable between different mice (Fig S6). *Irgm1*^−/−^ C-LP monocytes and Ly6C^+^MHC-II^+^ transitioning monocytes were the most severely affected, being outnumbered by WT cells >20 fold in some infected mice. That *C. rodentium* infection exacerbated the deficit of *Irgm1*^−/−^ myeloid cells in mixed hematopoietic chimera suggests that loss of Irgm1 severely compromised cellular fitness of C-LP myeloid cells during pathogen infection.

**Figure 6.**
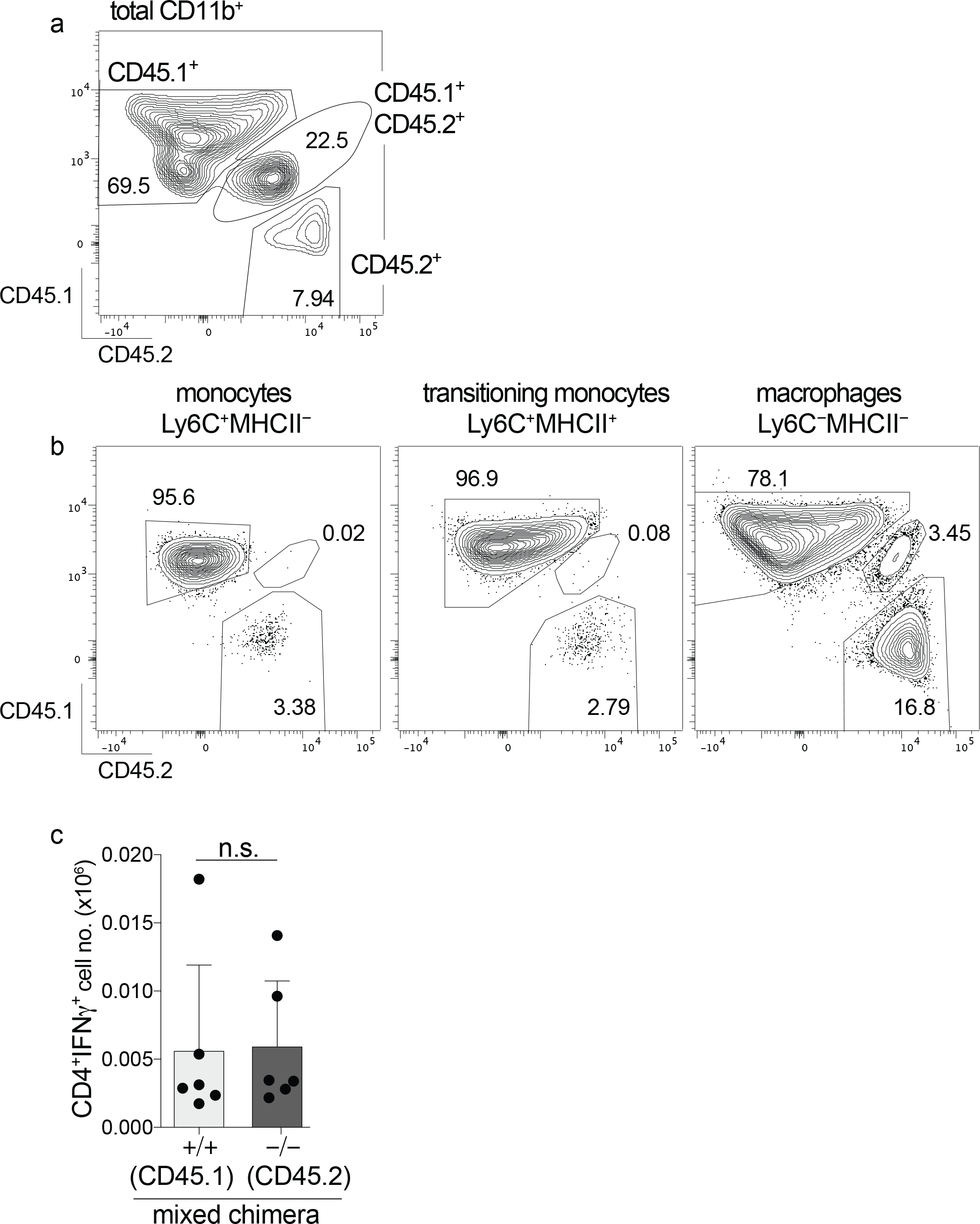
In mice harboring predominantly wild-type myeloid cells, Irgm1-deficient CD4 T cells mount a sufficient IFNγ response to *C. rodentium.* Mixed hematopoietic chimera reconstituted with a mixture of *Irgm1*^+/+^ (CD45.1) and *Irgm1*^−/−^ (CD45.2) bone marrow were analyzed for the relative abundance of C-LP immune cells originating from either of these genotypes, or from the host (WT, CD45.1^+^CD45.2^+^). (a-b) Representative flow plot of uninfected mixed hematopoietic chimera showing the relative abundance of host or donor-derived cells among total CD11b^+^ myeloid cells (a), or among the indicated subset of the monocyte/macrophage lineage (b). (c) Mixed hematopoietic chimera were infected for 10 days with *C. rodentium* and analyzed for the total number of IFNγ^+^ CD4 T cells originating from *Irgm1*^+/+^ or *Irgm1*^−/−^ bone marrow. Each dot represents one mouse. Error bars represent mean ± SD. n.s., not significant (unpaired Student’s t test).

Considering these results, we hypothesized that the decline of Irgm1-deficient myeloid cells in mixed hematopoietic chimera resulted from higher rates of cell death during *C. rodentium* infection. To address this question we used *Irgm1*^−/−^ mice, since all immune cell types were readily analyzed in these mice without the challenges imposed by low cell number and reduced competitiveness/fitness of Irgm1-deficient cells present in mixed chimera. We first analyzed cell death in monocytes and transitioning monocytes since these were the most robustly expanded populations during *C. rodentium* infection, and were populations in which Irgm1-deficiency had great impact on infection-induced remodeling (Fig 4). Indeed, the abundance of Annexin V^+^ C-LP monocytes in *C. rodentium* infected *Irgm1*^−/−^ mice was on average >20 fold higher than that of WT, with Annexin V expressed by as many as 40% of all C-LP monocytes (Fig 7a,b). By contrast, fewer than 1% of C-LP monocytes were Annexin V^+^ in infected WT mice, supporting the conclusions that *C. rodentium* infection was not inherently an apoptosis-inducing condition, and also, that increased monocyte apoptosis in *Irgm1*^−/−^ mice reflected a genotype-intrinsic susceptibility phenotype. Importantly, this susceptibility phenotype was induced by *C. rodentium* infection and was specific for monocytes within C-LP, since Annexin V expression was only marginally increased among bone marrow monocytes of *C. rodentium* infected mice, or among C-LP monocytes of uninfected mice (Fig S7). Also, neither of these monocyte populations had an increase in Annexin V^+^7AAD^+^ cells, which were significantly increased among C-LP monocytes in *C. rodentium* infected *Irgm1*^−/−^ mice (Fig 7b). To further delineate the requirements wherein Irgm1 function was key to prevent monocyte apoptosis within C-LP, we analyzed monocytes during DSS colitis. Following DSS treatment Annexin V expression on C-LP monocytes was similar between WT and *Irgm1*^−/−^ mice (Fig 7c,d), indicating that Irgm1 was not required to support monocyte survival in the setting of DSS colitis. Taken together, these studies suggest that *C. rodentium* infection is a unique stress in which Irgm1 has critical function to prevent dramatic rates of monocyte apoptosis.

**Figure 7.**
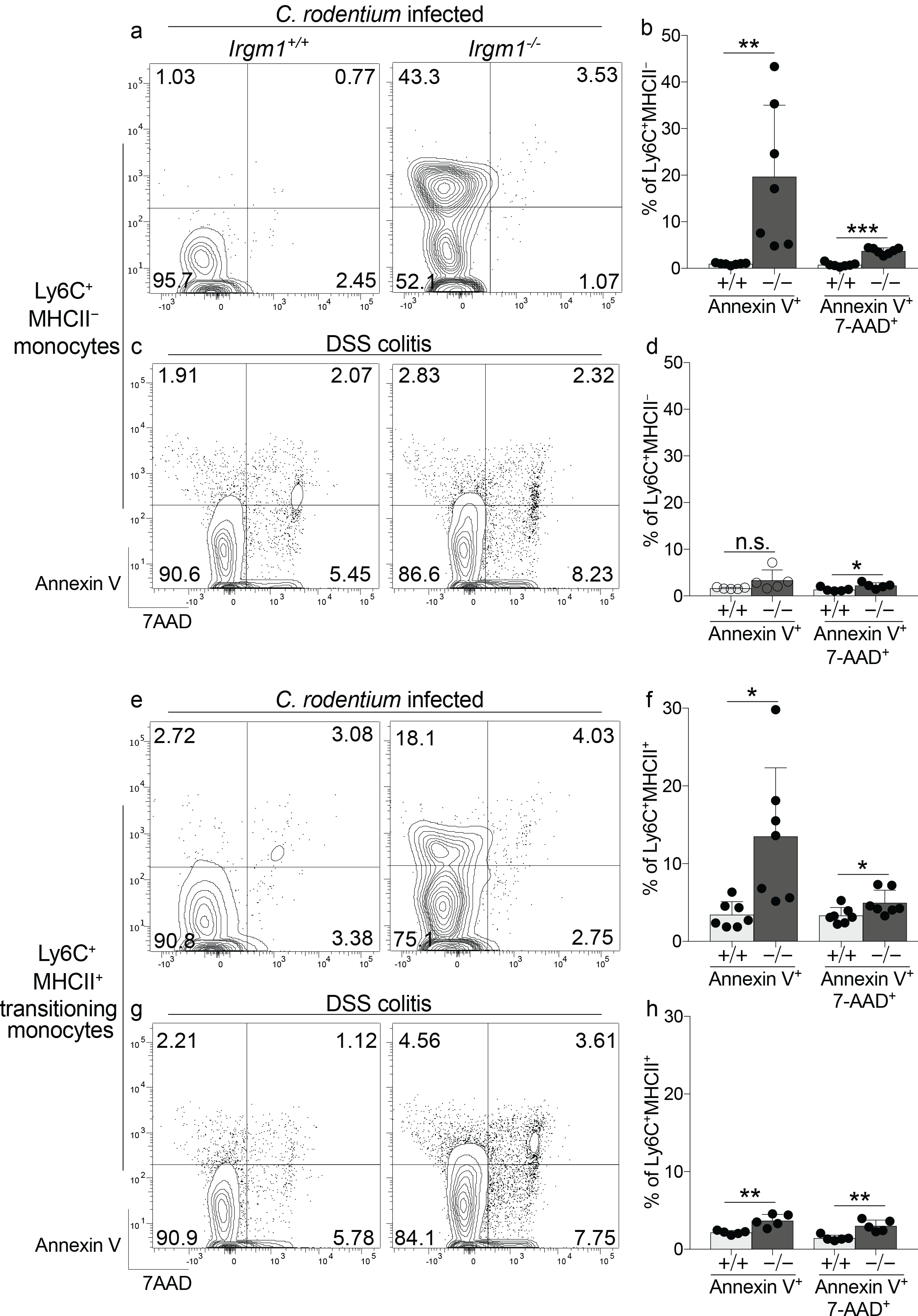
Specifically in the setting of *C. rodentium* infection, Irgm1 is required to prevent apoptosis of C-LP monocytes and transitioning monocytes. (a,b) Co-housed, littermate *Irgm1*^+/+^ and *Irgm1*^−/−^ mice were infected with *C. rodentium* and analyzed 10 days later for Annexin V and 7AAD expression on C-LP monocytes. Representative flow plot of C-LP monocytes from mice of the indicated genotype (a) and the percentage of Annexin V^+^ or Annexin V^+^7AAD^+^ monocytes from all mice tested (b). (c,d) C-LP monocytes from day 6 of DSS treated mice were analyzed as above. (e,f) *C. rodentium* infected mice from (a) were analyzed for Annexin V and 7AAD expression on C-LP transitioning monocytes. Representative flow plot of C-LP transitioning monocytes from mice of the indicated genotype (e) and the percentage of Annexin V^+^ or Annexin V^+^7AAD^+^ transitioning monocytes from all mice tested (f). (g,h) one mouse. Error bars represent mean ± \**p* < 0.05, \*\**p* < 0.01, \*\*\**p* < 0.001, n.s., not significant (unpaired Student’s t test).

Irgm1 function also prevented infection-induced apoptosis of C-LP Ly6C^+^MHCII^+^ transitioning monocytes, populations that are directly derived from monocytes and are typically robustly expanded in response to *C. rodentium* (Fig 7e,f). Infected *Irgm1*^−/−^ mice were markedly deficient in Ly6C^+^MHCII^+^ transitioning monocytes (Fig 4) and among the few Ly6C^+^MHCII^+^ cells that were present, Annexin V expression was >3 fold higher that of WT mice (Fig 7f). Similar to monocytes, the requirement for Irgm1 to prevent apoptosis of Ly6C^+^MHCII^+^ transitioning monocytes was both specific to and induced by *C. rodentium* infection, since there was little to no difference in apoptosis of these cells in uninfected mice, or in DSS-treated mice (Fig 7g,h and Fig S8). Infection-induced apoptosis of both monocytes and their Ly6C^+^MHCII^+^ progeny in *Irgm1*^−/−^ mice provide a rational explanation for why Ly6C^+^MHCII^+^ transitioning monocytes failed to expand in response to *C. rodentium* infection. Since monocytes/transitioning monocytes are precursors of C-LP macrophages, infection-induced apoptosis of these macrophage precursors is also likely to explain why Irgm1 was required to expand C-LP macrophages in response to *C. rodentium* infection. Consistent with this hypothesis the apoptotic susceptibility of macrophages, while increased at baseline, was not significantly enhanced by *C. rodentium* infection (Fig. S9), suggesting that failure to expand C-LP macrophages for *C. rodentium* immunity was primarily due to marked deficiency in the total number and overall viability of C-LP macrophage precursors, factors which were highly Irgm1-dependent. Paralleling our findings on monocytes, *C. rodentium* infection-induced apoptosis was prominent in all subsets of C-LP DCs, and was specific to this setting since increased apoptosis was not evident in DSS colitis (Fig. S9). Taken together, our findings identify a novel Irgm1 function to prevent cellular apoptosis of C-LP mononuclear phagocytes, and reveal *C. rodentium* infection as a unique intestinal stress in which this Irgm1 function is critical. Cell death of mononuclear phagocytes and concomitant failure to remodel immune populations specialized for *C. rodentium* immunity is a key factor underpinning *C. rodentium* outgrowth and pathogen susceptibility in *Irgm1*^−/−^ mice.

## Discussion

Despite being identified in 2007 as a Crohn’s disease susceptibility gene, the role of *IRGM* – and the murine orthologue, *Irgm1* – in intestinal immunity has remained unclear. Prior functional studies have focused on macrophages, in which the absence of Irgm1/IRGM leads to altered autophagic function and decreased killing of intracellular bacteria. Current dogma presumes that the role for IRGM/Irgm1 in xenophagy is the primary function for the proteins in immunity, and it is certainly likely to be a pivotal component of antibacterial immunity in some contexts. However, the requirements for host defense are unique for each pathogen, and pathogen defense in the intestine is particularly unique, since an unbalanced immune response to pathogens, or to the resident microbiota, can lead to chronic intestinal inflammation. Here we show that Irgm1 is required for immunity to an intestinal pathogen, and delineate that this requirement depends on a novel function for Irgm1 to regulate survival of C-LP mononuclear phagocytes and to support infection-induced remodeling of C-LP immune populations specialized for immunity to the extracellular enteric bacterium *C. rodentium*.

At baseline, intestinal homeostasis is essentially normal in *Irgm1*^−/−^ mice and it is only upon infection with *C. rodentium* that multiple defects in C-LP immune cells become apparent, with marked deficiencies in the number of transitioning monocytes, macrophages, dendritic cells, and Th1 cells. Deficiencies in *C. rodentium*-induced Th1 responses may well be the cause for mortality in *Irgm1*^−/−^ mice, since Th1 responses are essential for *C. rodentium* clearance[26] and also because mortality and *C. rodentium* expansion in *Irgm1*^−/−^ mice occurred at timepoints where Th1 and other adaptive immune responses would be expected to play an essential role in host defense. Based on our results with mixed hematopoietic chimera, Th1 cells expanded for *C. rodentium* immunity do not have an intrinsic requirement for Irgm1. Instead, our data suggest that deficiency in *C. rodentium*-induced Th1 cells in *Irgm1*^−/−^ mice is due to marked deficits in the number of C-LP transitioning monocytes, macrophages, and DCs, populations which prime intestinal Th1 cells directly and/or support the expansion of *C. rodentium*-induced Th1 cells[34, 37, 38, 40, 41]. Monocytes/macrophages are thought to support Th1 immunity to *C. rodentium* by serving as the key source for IL-12[34]. Our data support this specialized function for C-LP monocytes/macrophages, and further suggest that the low levels of IL-12 in the colon tissues of *C. rodentium* infected *Irgm1*^−/−^ mice are linked to poor survival and low cell number of monocytes/macrophages in these mice. These deficits were also evident in DCs, and while Irgm1 was required to support the population abundance of all DC subsets during *C. rodentium* infection, the impact was distinct for each subset and our data suggest that Irgm1’s function in DCs may be most critical in those subsets that support Th1 immunity. Intestinal Th1 immunity to *Toxoplasma gondii* requires C-LP CD103^+^CD11b^−^ DCs [43], and it is likely that this DC subset is also critical for *C. rodentium* immunity, considering that CD103^+^CD11b^−^ DCs are the most abundant DC subset in the colon, and we show here that their population abundance during *C. rodentium* infection is highly Irgm1-dependent. Th17 immunity to *C. rodentium* has been linked to CD103^+^CD11b^+^ DCs[36], and so it is interesting that the deficit in CD103^+^CD11b^+^ DCs in infected *Irgm1*^−/−^ mice did not result in a corresponding decrease in Th17 responses. It is possible that the DCs remaining in *Irgm1*^−/−^ mice were sufficient for Th17 immunity, or that APCs other than conventional CD103^+^ DCs play a more prominent role in Th17 responses to *C. rodentium*[44]. Whatever the cause, it is clear that Th17 and Th1 responses have distinct requirements for Irgm1. Since monocytes/macrophages and DCs are both linked to Th1 immunity to *C. rodentium*, whether Irgm1 dysfunction in DCs alone, monocytes/macrophages alone, or the combined defects of both cell types underpins the failure to expand Th1 cells for *C. rodentium* immunity awaits further experiments with lineage-specific knockout mice once they are developed.

Our studies describe a novel function for Irgm1 to regulate apoptosis in multiple cell types, including monocytes and macrophages. Although previous studies concluded that Irgm1 deficiency had no negative impact on macrophage apoptosis[20], our studies on macrophage apoptosis deviate from prior work in several ways. First, while most work on Irgm1 has been conducted on bone marrow macrophages tested *in vitro*, we investigated primary cells in their native tissue environment and in so doing, revealed a novel role for Irgm1 as a regulator of apoptosis in C-LP macrophages and their monocyte precursors. Second, we show that this function of Irgm1 is critical only in specific settings of stress, since Irgm1 was required for immune cell survival during *C. rodentium* infection, but was dispensable in DSS colitis. This finding is particularly outstanding, since DSS colitis and *C. rodentium* infection trigger similar histological and immunological changes in the intestine, and both stimulate the remodeling and expansion of C-LP monocytes/transitioning monocytes. That the requirements for Irgm1 for cell survival are distinct in these two inflammatory settings highlights the unique cellular and molecular requirements that tailor C-LP immune responses and specialize them to confront different types of insults. The stimulus-specific requirements for Irgm1 also extend to intestinal epithelial cells, which require Irgm1 to express Reg3Ɣ in DSS colitis [17], yet during *C. rodentium* infection upregulate Reg3Ɣ in an Irgm1-independent fashion. Potential explanations as to why Irgm1’s function to prevent monocyte apoptosis is most critical during *C. rodentium* infection may be that pathogen-specific factors trigger apoptosis directly[45, 46], or that these factors induce specific changes in C-LP monocytes/macrophages, which reasonably have distinct molecular requirements based on the nature of the stimulus. The specific mechanism through which lack of Irgm1 leads to apoptosis is currently unclear, but may relate to previous roles defined for Irgm1/IRGM in regulating mitochondrial dynamics and autophagy, both of which are heavily intertwined with apoptosis[47, 48]. This will be a key area in which to focus future research.

In summary, our results expand the paradigm for Irgm1/IRGM-mediated host defense by demonstrating a critical requirement for Irgm1/IRGM proteins to control the survival and remodeling of monocytes and macrophages that expand during enteric infection and defend against inflammatory bacteria. Failure to remodel and/or sustain immune cells which are specialized to manage such bacteria represent a novel mechanism that may underpin CD pathogenesis linked to *IRGM* dysfunction.

## Materials and Methods

### Mice

*Irgm1*^−/−^ mice have been described previously [2, 19]. They have been backcrossed to C57Bl/6NCr1 mice for 9 generations and were maintained by breeding Irgm1^+/−^ mice. The *Irgm1*^+/+^ and *Irgm1*^−/−^ mice used in these experiments were littermates and were co-housed under specific pathogen free (SPF) conditions. All procedures were approved by the IACUC of the Durham VA and Duke University Medical Centers.

### Citrobacter rodentium infection

Mice were infected by oral gavage with 0.1 ml of an overnight culture of Luria broth (LB) containing ~2.5 × 10^8^ colony-forming units of streptomycin-resistant *C. rodentium* (formerly *C. freundii* biotype 4280, strain DBS100). Mice were weighed daily and monitored for signs of illness or distress. Bacterial counts were determined in fresh catch fecal pellets or sterilely-excised spleens by homogenization in PBS, followed by plating of serial dilutions of the homogenate on LB/streptomycin plates, with colony counts assessed the following day.

### Histopathological scoring

Cecum and distal colon tissue from day 10 of *C. rodentium* infected mice were fixed in formalin and paraffin sections were stained using hematoxylin and eosin. Blinded histological scores were assigned using a previously validated scale[49]. The tissue was assessed for: submucosal edema (0 = no change; 1 = mild; 2 = moderate; and 3 = profound), epithelial hyperplasia (scored based on percentage greater than thickness of control epithelium, where 0 = no change; 1 = 1-50%; 2 = 51-100%; and 3 =>100%), epithelial integrity (0 = no change; 1 = <10 epithelial cells shedding per lesion; 2 = 11-20 epithelial cells shedding per lesion; 3 = epithelial ulceration; and 4 = epithelial ulceration with severe crypt destruction) and neutrophil and mononuclear cell infiltration (0 = none; 1 = mild; 2 = moderate; and 3 = severe), as previously described. The composite score was obtained by adding together the four individual scores for a maximum score of 13 points.

### Quantitative RT-PCR

Total RNA was extracted from colon tissue using an RNeasy Mini Kit (Qiagen, Valencia, CA) per the manufacturer’s instructions. Complementary DNA was generated using SuperScript II reverse transcriptase (Invitrogen, Carlsbad, CA) or an iScript cDNA Synthesis Kit (Bio-Rad). Quantitative RT-PCR was performed for each sample in duplicate, using TaqMan Gene Expression Master Mix (Applied Biosystems, Pleasanton, CA) or SsoAdvanced Universal SYBR Green Supermix (Bio-Rad). The following primer sets were used:

*Tnfα* forward 5’-ACCCTCACACTCAGATCATCTTCTC-3’, reverse 5’-TGAGATCCATGCCGTTGG-3’;
*Il1β* forward 5’-GCCACCTTTTGACAGTGATGAG-3’, reverse 5’-TGCTGCGAGATTTGAAGCTG-3’;
*Il12a* (p35) forward 5’-CTGTGCCTTGGTAGCATCTATG-3’, reverse 5’-GCAGAGTCTCGCCATTATGATTC-3’[34];
*Il12b* (p40) forward 5’-TGGTTTGCCATCGTTTTGCTG-3’ reverse 5’-ACAGGTGAGGTTCACTGTTTCT-3’;
*Il23a* (p19) forward 5’-CCAGCGGGACATATGAATCTACTAA-3’, reverse 5’-TTTGCAAGCAGAACTGGCTGT-3’
*Il22* forward 5’-CCGAGGAGTCAGTGCTAAGG-3’ reverse 5’-CATGTAGGGCTGGAACCTGT-3’;
*Actb* forward 5’-CGGTTCCGATGCCCTGAGGCTCTT-3’ reverse 5’-CGTCACACTTCATGATGGAATTGA-3’[50];
*Reg3Ɣ* from Applied Biosystems (Mm00441127_m1).

Relative gene expression was determined using the comparative threshold cycle (Ct) method[51]. Briefly, fold change was calculated using the equation, where ΔΔCt = [(Ct gene of interest - Ct actb)_Irgm1 KO_ - mean(Ct gene of interest - Ct actb)_Control WT_]. *Actb* was used as an internal control. The control WT group (non-infected) was set at a baseline of “1” by dividing all final values by the average of this group.

### Bacterial survival assays in macrophages

Primary murine bone marrow macrophage (BMM) cultures were established by flushing marrow from tibia and femurs using a 27g syringe filled with Dulbecco’s modified Eagle Medium (DMEM) (Life Technologies, Gaithersburg, MD); the marrow was then dispersed by drawing through the syringe a few times, and red cells were lysed with ammonium chloride. Adherent cells were cultured in BMM medium (DMEM supplemented with 10 % (v/v) fetal bovine serum (FBS) (Hyclone, Logan, UT) and 30 % (v/v) L929 cell-conditioned medium) for 5-6 days. For bacterial survival and immunostaining assays, the cells were lifted using Cell Dissociation Medium (Life Technologies), replated at equal densities, and then cultured for an additional 48 h before the assay. BMM were plated in 24-well plates at 0.35 × 10^6^ cells per well. They were exposed to 100u/mL IFN-γ (EMD Biosciences/ Calbiochem) for 24 h to activate them and induce Irgm1 expression. The cells were infected with *C. rodentium* at a multiplicity of infection (moi) of 2 bacteria per macrophage. The bacteria were added as a suspension in DMEM; the plates were centrifuged at 250 × g for 5 min, incubated for an additional 10 min, and washed 3 times with DMEM to remove extracellular bacteria. The cells were then incubated for various times in BMM medium supplemented with 6 μg/mL gentamycin. To quantify bacterial levels, the cells were washed with DMEM three times, and then lysed with 0.2 % (v/v) triton X-100 in PBS. Dilutions of the lysates were plated on LB plates, and colony forming units were counted. The assay was performed in triplicate for each condition.

### Preparation of colonic lamina propria and immune cell phenotyping

Colon lamina propria were processed as described[39] Briefly, colon was cut longitudinally and washed thoroughly to remove luminal contents. Colon was then cut into 0.3cm pieces and washed twice for 10 minutes in HBSS/10mM HEPES/0.035% NaHCO_3_/5mM EDTA/1.25% BSA/1mM DTT, followed by another 10 min wash in HBSS/10mM HEPES/0.035% NaHCO_3_/1.25%BSA. Tissue was transferred to C-tubes (Miltenyi) and digested using Liberase™ (Roche, 57.6 μg/ml) in the presence of DNase I (Sigma-Aldrich, 8 U/ml). Tissues were incubated with agitation for 30 min at 37°C, then dissociated using the GentleMACS™ Dissociator (Miltenyi). The resulting cellular suspension was filtered through a 100μm mesh (BD Falcon), centrifuged, and subjected to RBC lysis. After RBC lysis, cell suspension was filtered through a 40μm mesh (BD Falcon).

Freshly isolated lamina propria cells were stained with the following antibodies and reagents: FcR block (Biolegend), Live/dead® fixable dead cell staining (Thermo Fisher), anti-CD11c (N418), anti-IA/IE (M5/114.15.2), anti-CD11b (M1/70), anti-CD14 (Sa14-2), anti-CD24 (M1/69), anti-CD103 (2E7), anti-Ly6C (HK1.4), anti-Ly6G (1A8),anti-CD4 (GK1.5), anti-CD8a (53-6.7), anti-CD45 (30-F11), anti-TCRβ (H57-597), anti-IFNγ (XMG1.2), anti-IL-17A (TC11-18H10.1), anti-Foxp3 (FJK-16S), and in some cases, anti-CD45.2 (104) and anti-CD45.1 (A20). Some experiments also included Annexin V (Biolegend) and 7aad (Biolegend). For flow cytometry analysis, we gated populations from live cells according to the following markers: CD4 T cells, (CD45^+^TCRβ^+^CD4^+^); CD8 T cells, (CD45^+^TCRβ^+^CD8a^+^); DCs, (IA/IE^+^CD11c^+^CD24^+^CD14^−^, followed by CD103 and CD11b to distinguish DC subsets); Monocytes, (CD11b^+^Ly6C^+^CD24^−^IA/E^−^); Transitioning monocytes, (CD11b^+^Ly6C^+^IA/E^+^CD24^−^); Macrophages, (CD11b^+^IA/E^+^CD24^−^ Ly6C^−^); and Neutrophils, (CD11b^+^Ly6G^+^). Foxp3 staining was performed using Foxp3/Transcription Factor Staining Buffer Set (eBioscience). Annexin V staining and analysis was performed using Annexin V binding buffer (Biolegend). For intracellular cytokine staining, lamina propria cells were resuspended in complete RPMI and stimulated for 4h with PMA (50 ng/ml) and ionomycin (500 ng/ml) in the presence of BD Golgi plug (BD Biosciences), and then stained for intracellular cytokines using the Cytofix/Cytoperm kit (BD biosciences). Flow cytometry analysis of cells was performed on a BD Canto (BD Biosciences), and data further analyzed using FlowJo software.

### Hematopoietic chimera

To generate hematopoietic chimera, 10-week-old CD45.1^+^ WT mice were irradiated with two doses of 600 cGy (X-RAD 320), with 3h rest between doses. Irradiated mice were injected intravenously with 8×10^6^ bone marrow cells from CD45.2^+^ WT mice or from CD45.2^+^ *Irgm1*^−/−^ mice. To generate mixed hematopoietic chimera, 10-week-old CD45.1^+^CD45.2^+^ WT mice (bred in house) were irradiated as above and reconstituted with a total number of 9.6×10^6^ bone marrow cells composed of a mixture of bone marrow from WT (CD45.1) and *Irgm1*^−/−^ mice (CD45.2) at a 1: 5 ratio. All hematopoietic chimera were analyzed and/or subjected to *C. rodentium* infection 8 weeks following immune reconstitution.

### Dextran Sodium Sulfate (DSS)-induced colitis

Acute colitis was established according to standard protocols [52, 53] by adding 3% (w/v) dextran sodium sulfate (DSS) (ICN Biomedicals Inc., Aurora, Ohio) to the drinking water of mice for 6 days. Mice were analyzed on the last day of DSS treatment. Control mice received water without DSS. Body weights were recorded daily.

### Statistical analyses

Statistical comparisons were completed using Excel (Microsoft) or GraphPad Prism software (GraphPad, San Diego, CA) to perform 2-tailed Student’s t-test.

## Supplementary Figure Legends

**Supplementary Figure 1.**
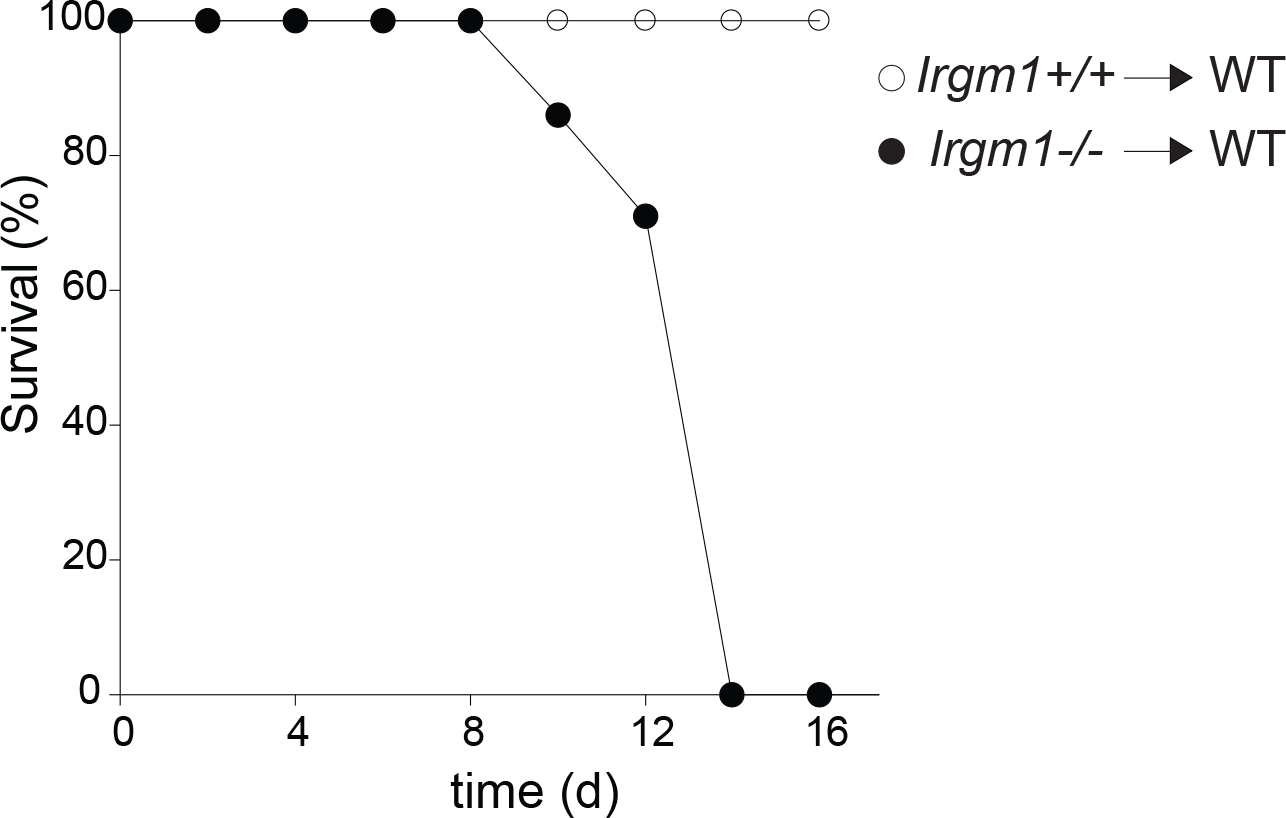
Survival of hematopoietic chimera reconstituted with *Irgm1*^+/+^ or *Irgm1*^−/−^ bone marrow following infection with *C. rodentium*. n=7. The difference in survival was statistically significant [p=0.0004, Log Rank (Mantel-Cox) test].

**Supplementary Figure 2.**
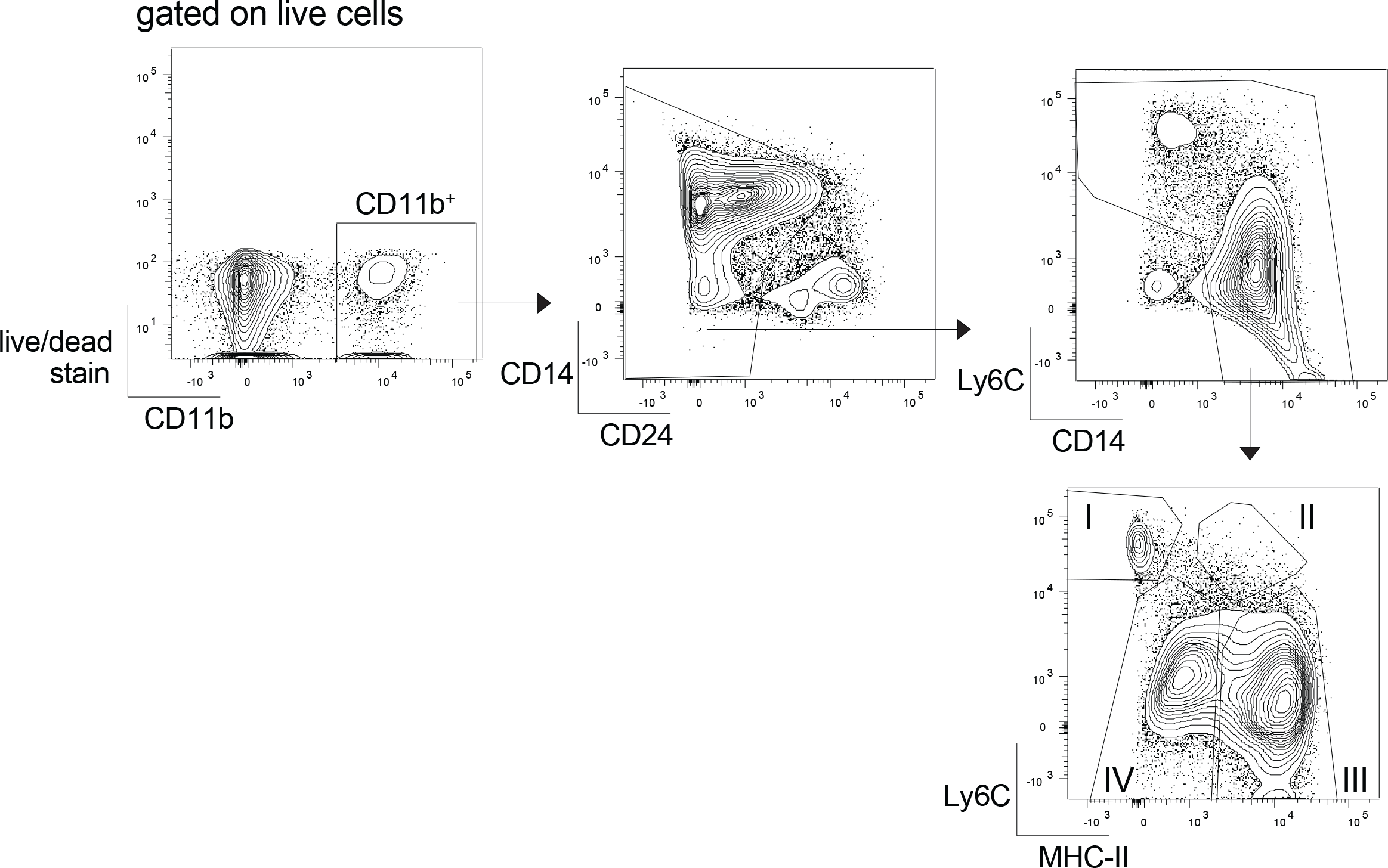
Gating strategy to analyze C-LP cells of the monocyte/macrophage lineage. After applying the indicated gates, populations were distinguished as follows: I. Monocyte (Ly6C^+^MHC-II^−^), II. transitioning monocyte (Ly6C^+^MHC-II^+^), III. macrophage (Ly6C^−^MHC-II^+^); IV. MHC-II^lo^ macrophage (Ly6C^−^MHC-II^lo^).

**Supplementary Figure 3.**
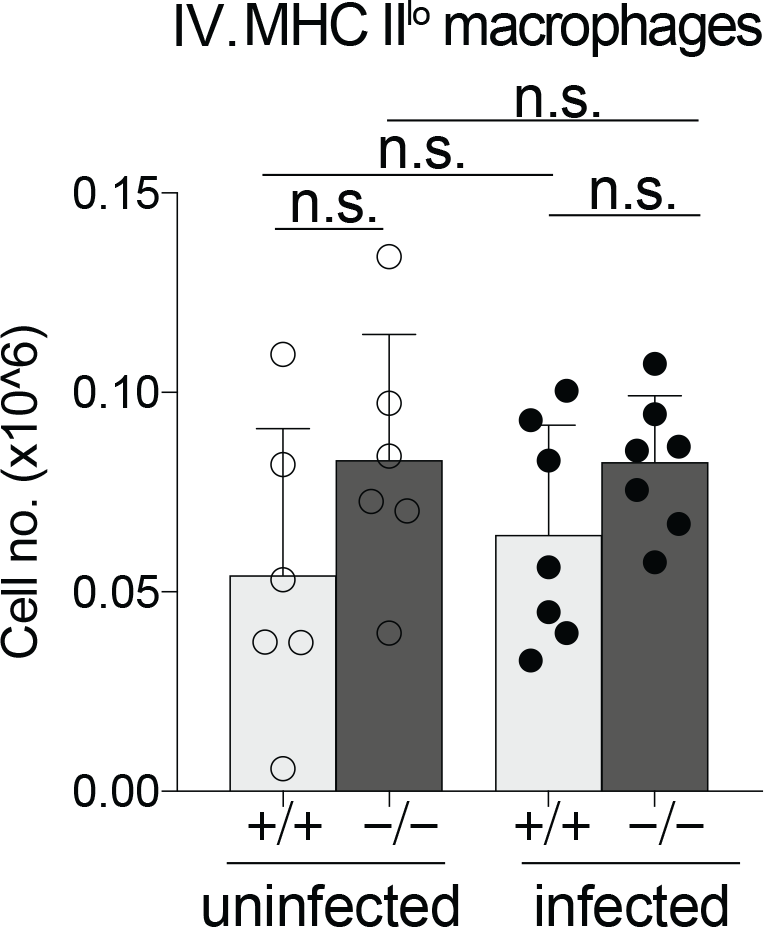
Total cell number of C-LP MHC-II^lo^ macrophages in uninfected or *C. rodentium* infected mice of the indicated genotype. Data were combined from two independent experiments. Error bars represent mean ± SD. n.s., not significant (unpaired Student’s t test).

**Supplementary Figure 4.**
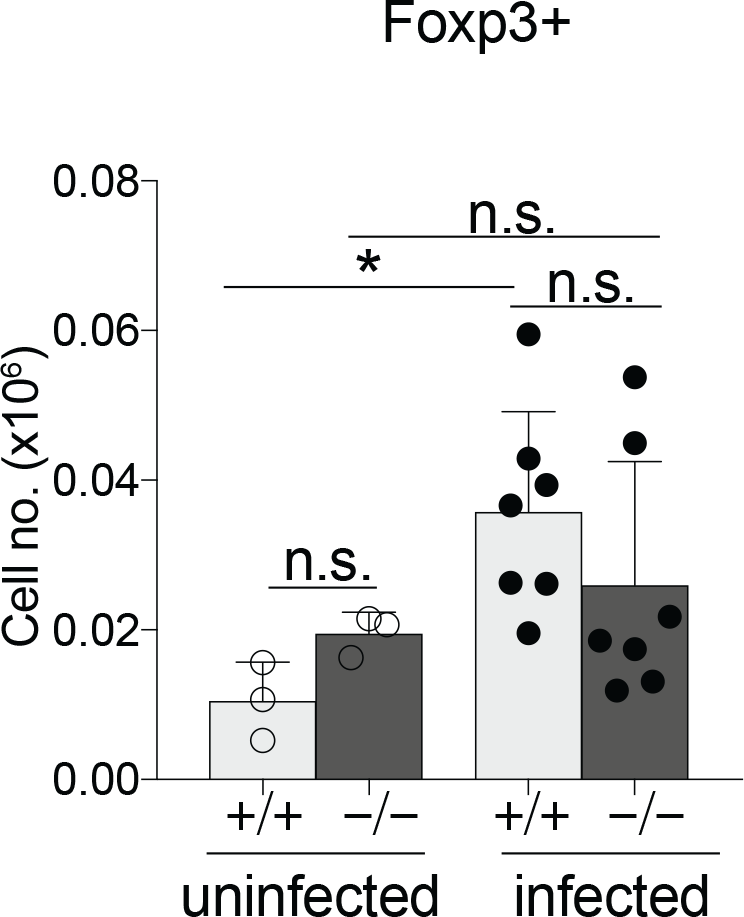
Total cell number of C-LP Foxp3+ CD4 T cells in uninfected or *C. rodentium* infected mice of the indicated genotype. Data were combined from two independent experiments. Error bars represent mean ± SD. \**p* < 0.05, n.s., not significant (unpaired Student’s t test).

**Supplementary Figure 5.**
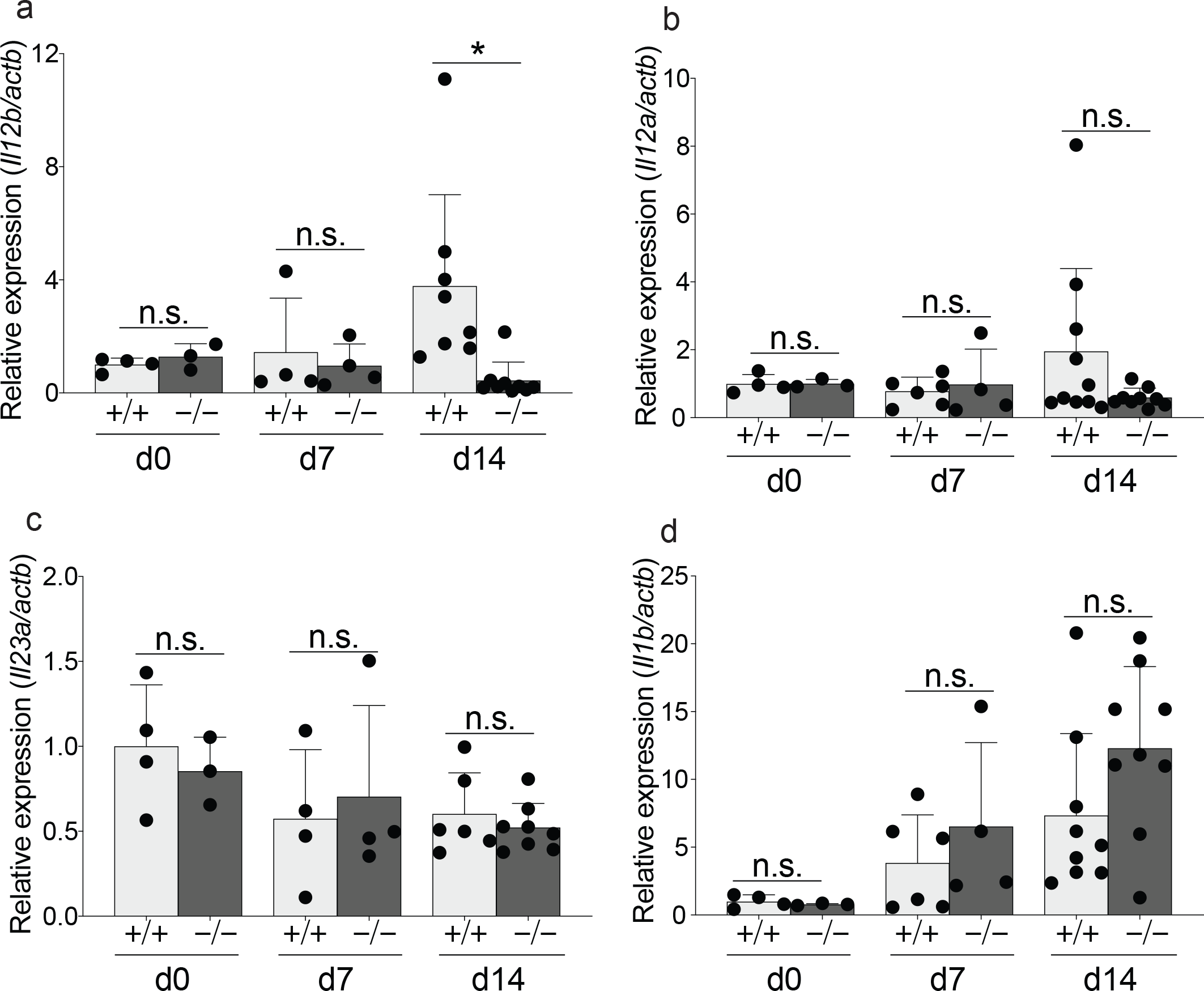
Distal colon from uninfected or *C. rodentium* infected mice described in Fig. 2e-g were analyzed for tissue expression of mRNA for IL-12b/p40 (a), IL-12a/p35 (b), IL-23/p19 (c), and IL-1β (d). Error bars represent mean ± SD. \**p* < 0.05, n.s., not significant (unpaired Student’s t test).

**Supplementary Figure 6.**
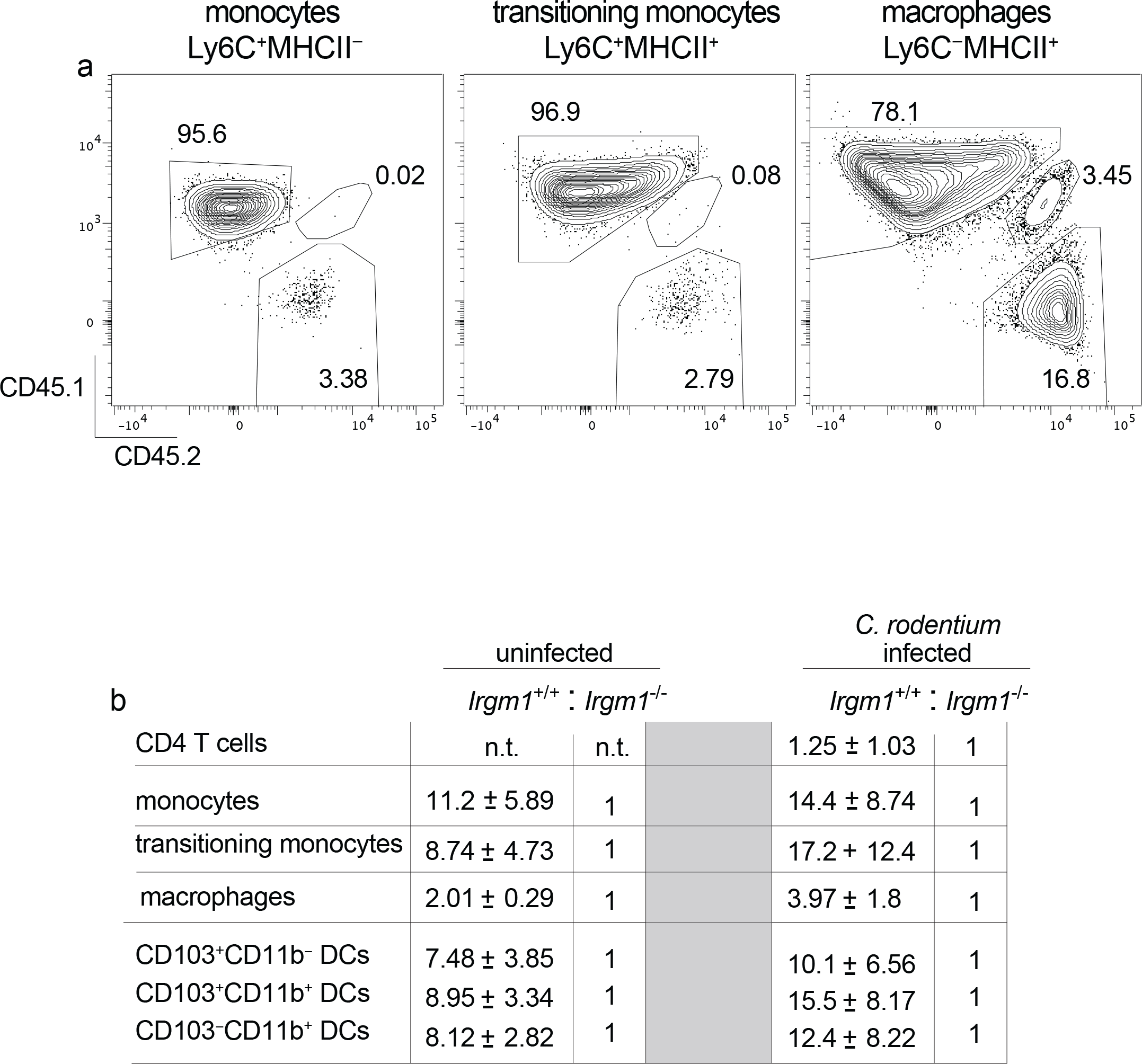
(a-b) Mixed hematopoietic chimera described in Fig. 6 were infected with *C. rodentium* for 10 days and analyzed for C-LP immune cells originating from *Irgm1*^+/+^ (CD45.1), *Irgm1*^−/−^ (CD45.2), or host-derived (CD45.1^+^CD45.2^+^) cells. (a) Representative flow plots showing the relative abundance of host or donor derived cells among C-LP monocytes, transitioning monocytes, and macrophages in mixed hematopoietic chimera infected with *C. rodentium*. (b) The ratio of *Irgm1*^+/+^ to *Irgm1*^−/−^ cells among the indicated cell type in C-LP of mixed hematopoietic chimera, either uninfected or infected for 10 days with *C. rodentium*. Data represents mean ± SD. n.t., not tested.

**Supplementary Figure 7.**
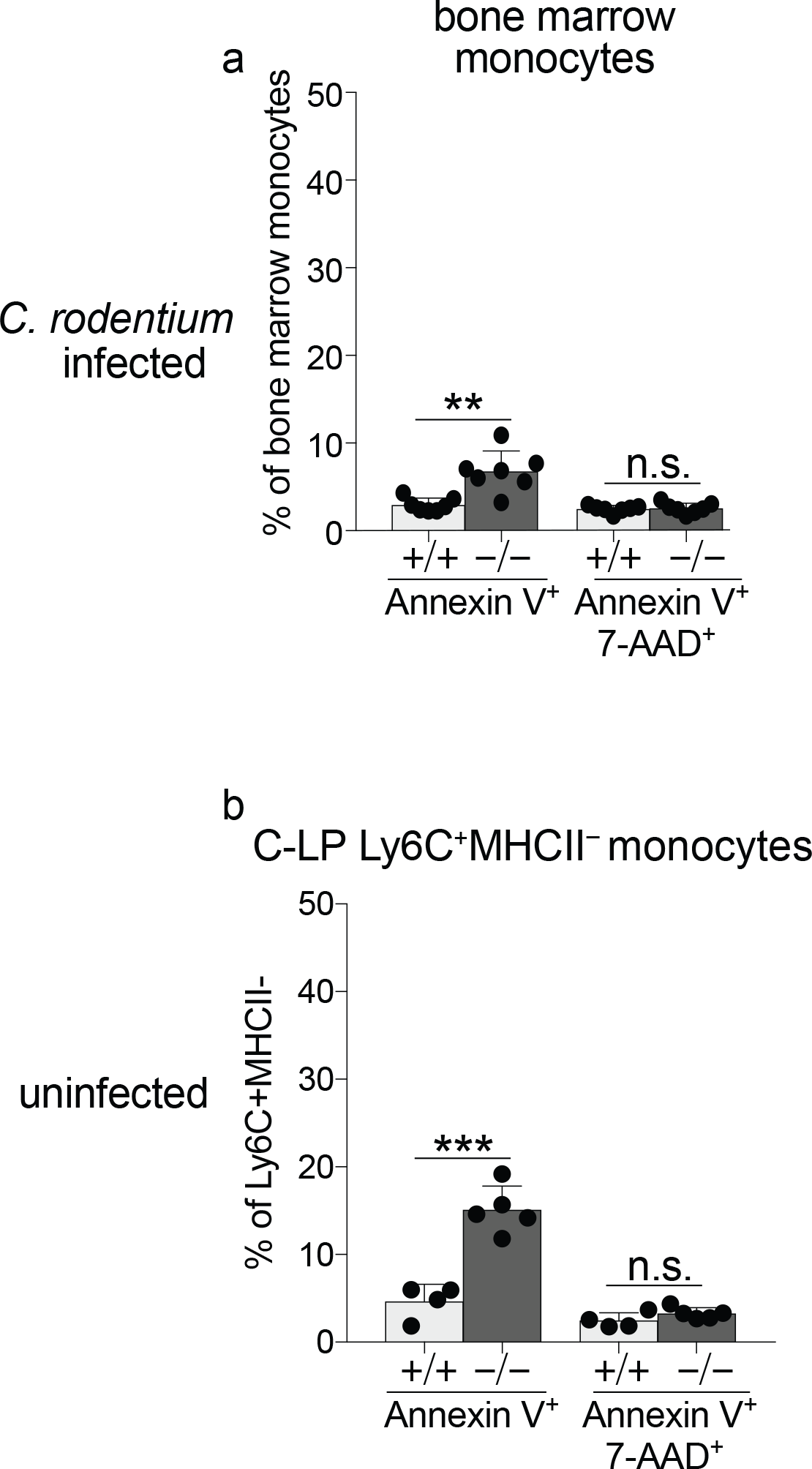
(a) *C. rodentium* infected mice described in Fig. 7 were analyzed for the percentage of Annexin V^+^ or Annexin V^+^7AAD^+^ cells among bone marrow monocytes. (b) C-LP monocytes from uninfected mice of the indicated genotype were analyzed as above. Each dot represents one mouse. Error bars represent mean ± SD. \*\**p* < 0.01, \*\*\**p* < 0.001, n.s., not significant (unpaired Student’s t test).

**Supplementary Figure 8.**
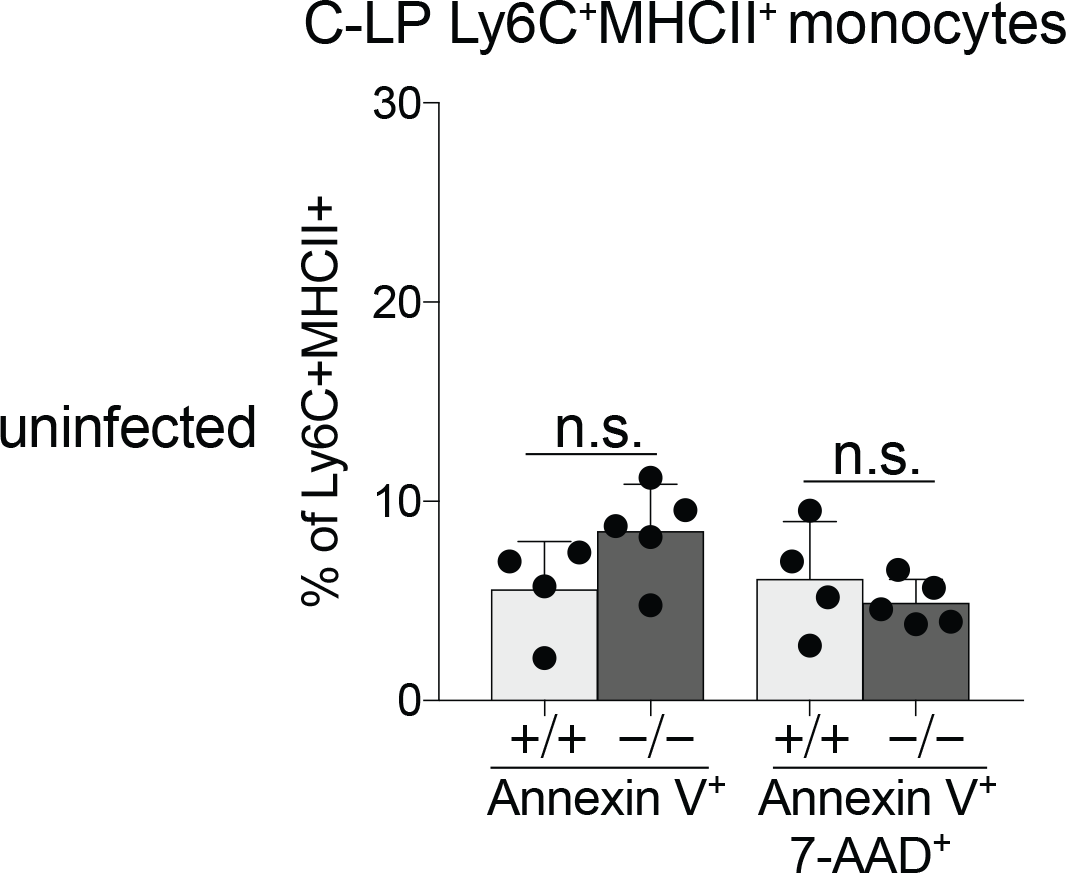
The percentage of Annexin V^+^ or Annexin V^+^7AAD^+^ cells among C-LP transitioning monocytes in uninfected mice. Each dot represents one mouse. Error bars represent mean ± SD. n.s., not significant (unpaired Student’s t test).

**Supplementary Figure 9.**
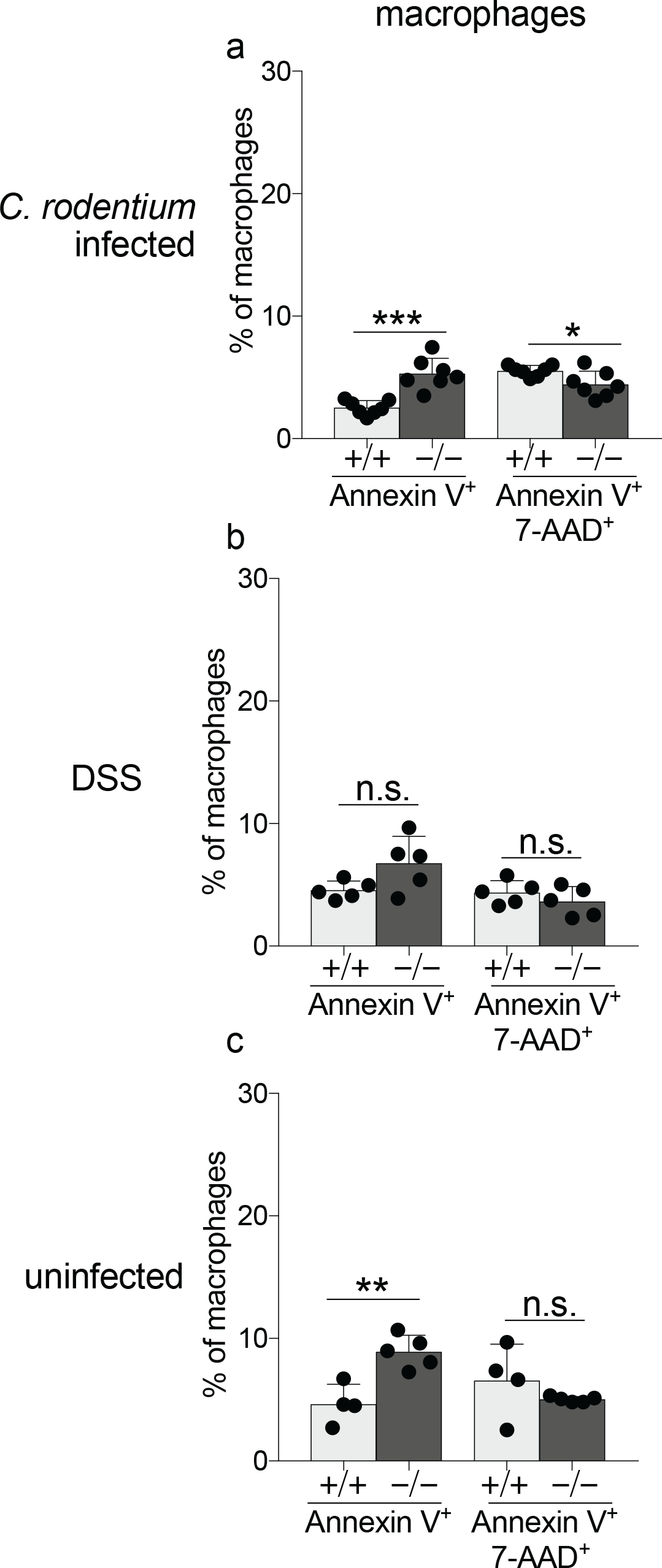
The percentage of Annexin V^+^ or Annexin V^+^7AAD^+^ cells among MHC-II^hi^ C-LP macrophages among *C. rodentium* infected (a) DSS treated (b) or uninfected mice (c) of the indicated genotype. Each dot represents one mouse. Error bars represent mean ± SD. \**p* < 0.05, \*\**p* < 0.01, \*\*\**p* < 0.001, n.s., not significant (unpaired Student’s t test).

**Supplementary Figure 10.**
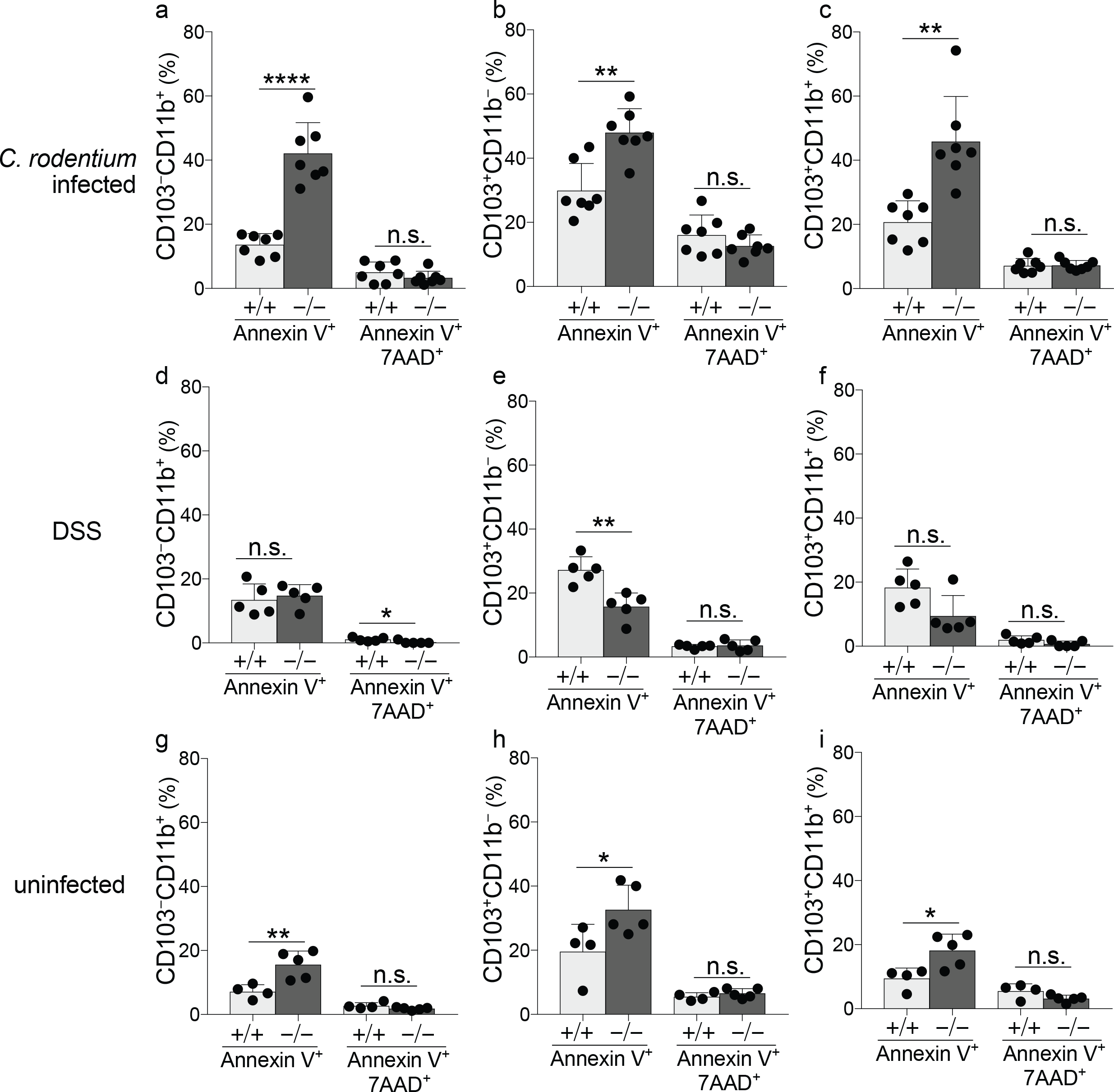
The percentage of Annexin V^+^ or Annexin V^+^7AAD^+^ cells among C-LP CD103^−^CD11b^+^ DCs (a), CD103^+^CD11b^−^ DCs (b), and CD103^+^CD11b^+^ DCs (c) in *C. rodentium* infected mice described in Fig 7. C-LP DCs from DSS treated mice (d-f) or uninfected/untreated mice (g-i) were analyzed as above. Error bars represent mean ± SD. \**p* < 0.05, \*\**p* < 0.01, \*\*\*\**p* < 0.0001, n.s., not significant (unpaired Student’s t test). SD. \**p* < 0.05, \*\**p* < 0.01, \*\*\*\**p* < 0.0001 (unpaired Student’s t test).

## Acknowledgments

B.A.V. is the Children with Intestinal and Liver Disorders (CH.I.L.D.) Foundation Chair in Pediatric Gastroenterology.

